# SVEP1, a novel human coronary artery disease locus, promotes atherosclerosis

**DOI:** 10.1101/2020.06.15.151027

**Authors:** In-Hyuk Jung, Jared S. Elenbaas, Arturo Alisio, Katherine Santana, Erica P. Young, Chul Joo Kang, Puja Kachroo, Kory J. Lavine, Babak Razani, Robert P. Mecham, Nathan O. Stitziel

## Abstract

A low-frequency variant of SVEP1, an extracellular matrix protein, is associated with risk of coronary disease in humans independent of plasma lipids. Despite a robust statistical association, however, it was unclear if and how SVEP1 might contribute to atherosclerosis. Here, using Mendelian randomization and complementary mouse models, we provide evidence that SVEP1 promotes atherosclerosis in humans and mice. We find that SVEP1 is expressed by vascular smooth muscle cells (VSMCs) within the atherosclerotic plaque. VSMCs also interact with SVEP1, causing proliferation and dysregulation of key differentiation pathways, including integrin and Notch signaling. Fibroblast growth factor receptor transcription increases in VSMCs interacting with SVEP1, and is further increased by the coronary disease-associated SVEP1 variant. These effects ultimately drive inflammation and promote atherosclerosis. Taken together, our results suggest that VSMC-derived SVEP1 is a pro-atherogenic factor, and support the concept that pharmacological inhibition of SVEP1 should protect against atherosclerosis in humans.

## Introduction

Cardiometabolic diseases are leading causes of morbidity and mortality and their prevalence is increasing (Dzau et al., 2002; Hansson and Klareskog, 2011; Liu and Ntambi, 2009; Rader and FitzGerald, 1998; Randolph, 2013; Ross, 1996; Virella and Lopes-Virella, 2003; Weber and Noels, 2011). Although approved treatments can help ameliorate these diseases, residual disease risk remains a substantial problem. Statin medications, for example, lower plasma cholesterol levels and reduce risk of coronary events by 20-30% (C Baigent, 2005), highlighting both significant residual risk and an unmet need for identifying alternative treatment strategies. Human genetics is a powerful approach to uncover potential therapeutic targets and to date more than 160 loci (Deloukas et al., 2013; Nelson et al., 2017; Nikpay et al., 2015; Schunkert et al., 2011) have been robustly associated with coronary artery disease (CAD). At most loci, however, the causal gene is unknown, presenting a major bottle-neck and hindering the translation of these findings into new therapies. We previously performed a large-scale exome-wide association study of low-frequency protein altering variation and identified a highly conserved missense polymorphism in *SVEP1* (p.D2702G) that associated with an increased risk of CAD (Odds Ratio 1.14 per risk allele) (Stitziel et al., 2016). This CAD risk variant (hereto referred to as SVEP1^CADrv^) is not associated with an effect on plasma lipids but has a modest positive association with blood pressure and type 2 diabetes (Stitziel et al., 2016), suggesting this variant may broadly contribute to the progression of cardiometabolic disease.

*SVEP1*, also known as polydom, encodes a large extracellular matrix protein with sushi (complement control protein), von Willebrand factor type A, epidermal growth factor-like (EGF), and pentraxin domains (Gilges et al., 2000; Shur et al., 2006). This gene was originally discovered in a screen for Notch-interacting proteins, as it contains Notch-like repeat EGF-domains (Shur et al., 2006). The only protein currently known to directly interact with SVEP1 is integrin α9β1 (Sato-Nishiuchi et al., 2012), a provisional matrix-binding integrin that is also linked to increased blood pressure in humans(Levy et al., 2009; Takeuchi et al., 2010). Integrin α9β1 binds to the same protein domain that harbors the variant residue in SVEP1^CADrv^ (Sato-Nishiuchi et al., 2012) and both proteins also play critical roles in development, including lymphatic patterning (Karpanen et al., 2017; Samuelov et al., 2017).

Despite the strong statistical evidence linking *SVEP1* with CAD, its direct causality and associated mechanisms were unclear. Here, we sought to determine if and how SVEP1 may influence the development of atherosclerosis. Given the overlapping disease associations between SVEP1 and integrin α9β1, their shared biological functions, and the proximity of the variant to integrin α9β1’s binding site, we focused our mechanistic studies on cell types that play a prominent role in atherosclerosis and express SVEP1 and/or integrin α9β1.

## Results

### *SVEP1* is expressed by arterial VSMCs under pathological conditions

To begin characterizing the role of SVEP1 in the pathogenesis of atherosclerosis, we sought to identify disease-relevant tissues and cell types that express *SVEP1*. Expression data from the Genotype-Tissue Expression (GTEx) project indicate that human arterial tissue, including coronary arteries, express *SVEP1* (Figure S1A). To confirm arterial expression, we used *in situ* hybridization on tissue explants from the aortic wall and internal mammary artery of patients with established coronary artery disease. *SVEP1* expression was readily detected within cells staining with the vascular smooth muscle cell marker, smooth muscle α-actin (SMα-actin) (Figure 1A). VSMCs are known to increase synthesis of certain extracellular matrix proteins in response to various pathological stimuli (Cangemi et al., 2011); therefore, we assessed expression data from relevant disease specimens to determine if this also applies to SVEP1. Indeed, *SVEP1* expression is higher within human atherosclerotic tissue from carotid explants, relative to patient-paired adjacent and macroscopically intact tissue (Ayari and Bricca, 2013) (Figure S1B). Athero-prone arterial tissue explants from patients with diabetes also express higher levels of *SVEP1* compared to patients without diabetes (Cangemi et al., 2011) (Figure S1C).

**Figure 1.**
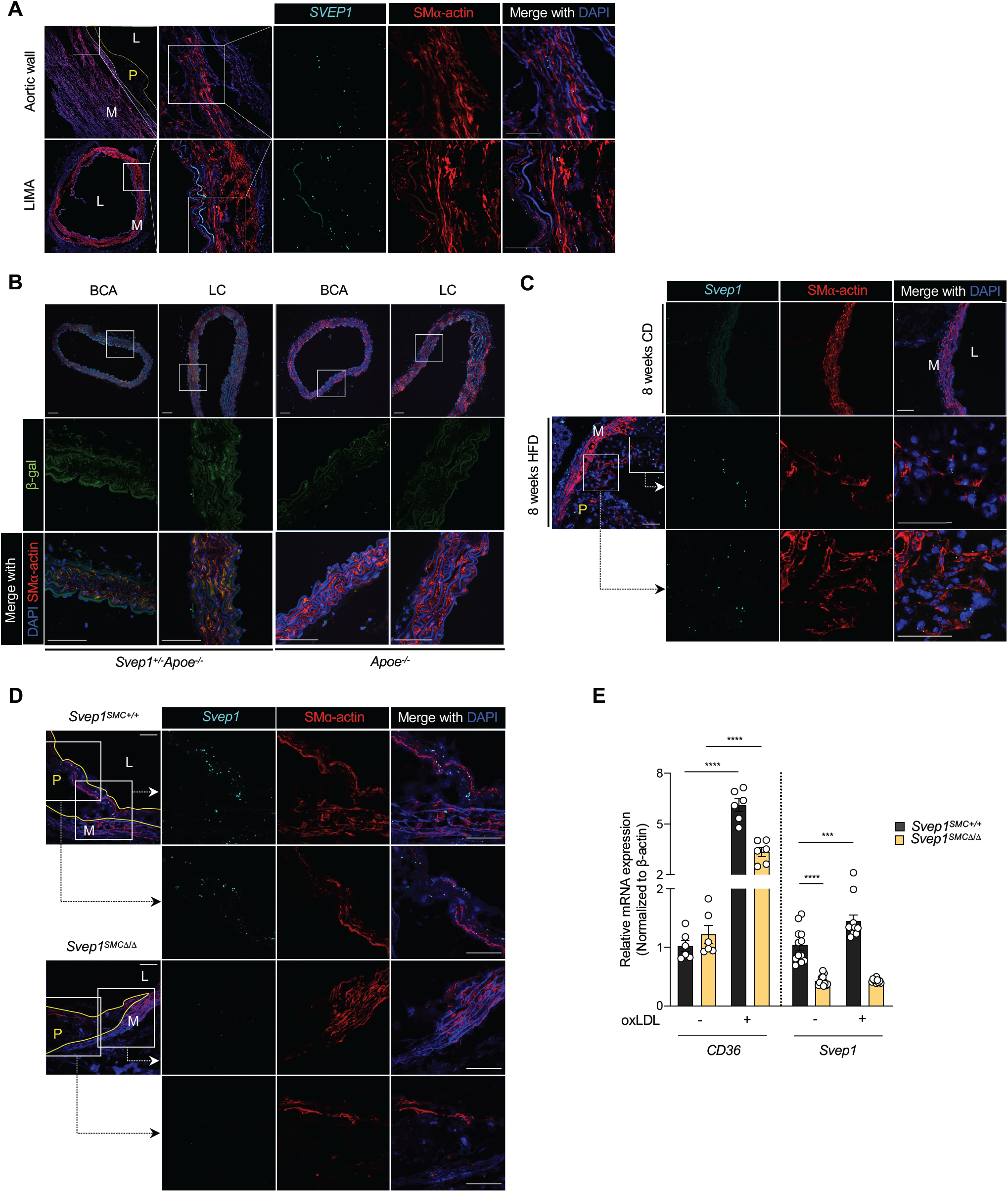
*SVEP1* is expressed by VSMCs under pathological conditions. (A) Expression of *SVEP1* in human aortic wall and LIMA cross-sections from patients using ISH. (B) β-gal expression in the aortic root, BCA (brachiocephalic artery), LC (lesser curvature) from 8-week-old *Svep1*^*+/-*^*Apoe*^*-/-*^ and *Apoe*^*-/-*^ mice. (C) Expression of *Svep1* using ISH in aortic root from young (8-week-old), CD fed and 8 weeks of HFD fed *Apoe*^*-/-*^ and *Svep1*^*+/-*^*Apoe*^*-/-*^ mice. (D) Expression of *Svep1* using ISH in the aortic root from *Svep1*^*SMC+/+*^ and *Svep1*^*SMCΔ/Δ*^ mice after 8 weeks of HFD feeding. Outlined areas indicate the regions magnified in the next panels. Tissues in (A-D) were co-stained with the VSMC marker, SMα-actin. Scale bars, 50 µm. M, media; L, lumen; P, plaque. (E) *Svep1* expression of primary VSMCs from *Svep1*^*SMC+/+*^ and *Svep1*^*SMCΔ/Δ*^ mice with or without the addition of oxLDL for 48 hr. Increased expression of CD36, the oxLDL receptor, confirms VSMC stimulation. \*\*\**P* < 0.001; \*\*\*\**P* < 0.0001. Mann-Whitney test was used.

We obtained mice expressing a lacZ reporter under the native *Svep1* promotor to determine whether murine *Svep1* expression recapitulated human *SVEP1* expression, and may therefore be a viable animal model to study its effects on disease. Within healthy arterial tissue of young mice, we observed low β-gal expression, mostly colocalizing with VSMCs (Figure 1B). These data are consistent with published single-cell studies that identify VSMCs within the healthy murine aorta as a minor source of *Svep1* expression (Kalluri et al., 2019) (Figure S1D). To determine if murine *Svep1* expression was increased in the development of atherosclerosis, as in humans, we assayed expression within mouse arterial tissue after the induction of atherosclerosis. Experimental atherosclerosis was induced by feeding atheroprone (*Apoe*^*-/-*^) mice a Western, high-fat diet (HFD) for 8 weeks. *Apoe*^*-/-*^ mice fed a standard chow diet (CD) served as non-atherogenic controls. After 8 weeks of an atherogenic HFD, we observed a 2-fold increase in *Svep1* expression relative to CD fed control mice (Figure 1C, S1E). This expression was colocalized with neointimal cells that co-stained with SMα-actin, suggesting VSMC expression (Figure 1C).

Numerous cell types have been demonstrated to gain expression of VSMC markers in the context of atherosclerosis (Bennett et al., 2016). Therefore, to test the hypothesis that VSMC-derived cells within the neointima are the predominate source of Svep1, we generated *Apoe*^*-/-*^ mice with VSMC-specific knockout of *Svep1* (*Svep1*^*flox/flox*^*Myh11-Cre*^*ERT2*^*Apoe*^*-/-*^; hereafter referred to as *Svep1*^*SMCΔ/Δ*^) and mice with unaltered *Svep1* expression (*Svep1*^*+/+*^*Myh11-Cre*^*ERT2*^*Apoe*^*-/-*^; hereafter referred to as *Svep1*^*SMC+/+*^), which served as controls. *Svep1* expression was assessed using *in situ* hybridization within the neointima of the aortic root of both groups after 8 weeks of HFD feeding. Indeed, while we observed robust *Svep1* expression in control mice, neointimal *Svep1* expression was nearly undetectable in *Svep1*^*SMCΔ/Δ*^ mice (Figure 1D, S1F). These data indicate that VSMC-derived cells are the major source of *Svep1* in atherosclerotic plaque.

Given the increased expression of *Svep1* under atherosclerotic conditions in mice and humans, we tested the ability of atheroma-associated oxidized LDL (oxLDL) to directly induce *Svep1* expression in VSMCs. Exposure to oxLDL increased *Svep1* expression by 48% in primary VSMCs from *Svep1*^*SMC+/+*^ mice but not *Svep1*^*SMCΔ/Δ*^ mice, compared to vehicle-treated control cells (Figure 1E). Both *Svep1*^*SMC+/+*^ and *Svep1*^*SMCΔ/Δ*^ cells increased expression of *CD36*, indicating they were activated upon binding of oxLDL with its receptor (Wei Li, 2010).

Taken together, these data demonstrate that *SVEP1* is produced locally by VSMCs in atherosclerotic disease and are consistent with prior studies suggesting SVEP1 is expressed by cells of mesenchymal origin (Karpanen et al., 2017; Morooka et al., 2017). Further, these data suggest that SVEP1 may play a direct role in the pathogenesis of atherosclerosis and that mouse models are an appropriate means to interrogate this question.

### *Svep1* drives atherosclerotic plaque development

To study the effect of *Svep1* on atherosclerosis, we fed *Apoe*^*-/-*^ and *Svep1*^*+/-*^*Apoe*^*-/-*^ mice (mice with homozygous *Svep1* deficiency die from edema at day E18.5 (Karpanen et al., 2017; Morooka et al., 2017) a HFD for 8 weeks and analyzed the resulting atherosclerotic plaque burden. There were no observed differences between genotypes in body weight, plasma total cholesterol, triglycerides, and glucose (Figure 2A, B). Relative to controls, however, *Svep1*^*+/-*^*Apoe*^*-/-*^ mice had a significant reduction in plaque burden (as characterized by the percentage of surface area staining positive with Oil Red O) in the aortic arch and whole aorta by *en face* preparations, as well as in sectioned aortic roots (Figure 2C, D). *Svep1* deficiency also resulted in reduced macrophage staining within the aortic root neointima, as determined by the percentage of area staining positive for Mac3 (Figure 2E). We did not appreciate marked differences in measures of plaque stability, such as area staining positive for VSMC markers or necrotic core size, although collagen content was modestly higher in atheromas from control mice compared to *Svep1*^*+/-*^*Apoe*^*-/-*^ mice (Figure S2A-C).

**Figure 2.**
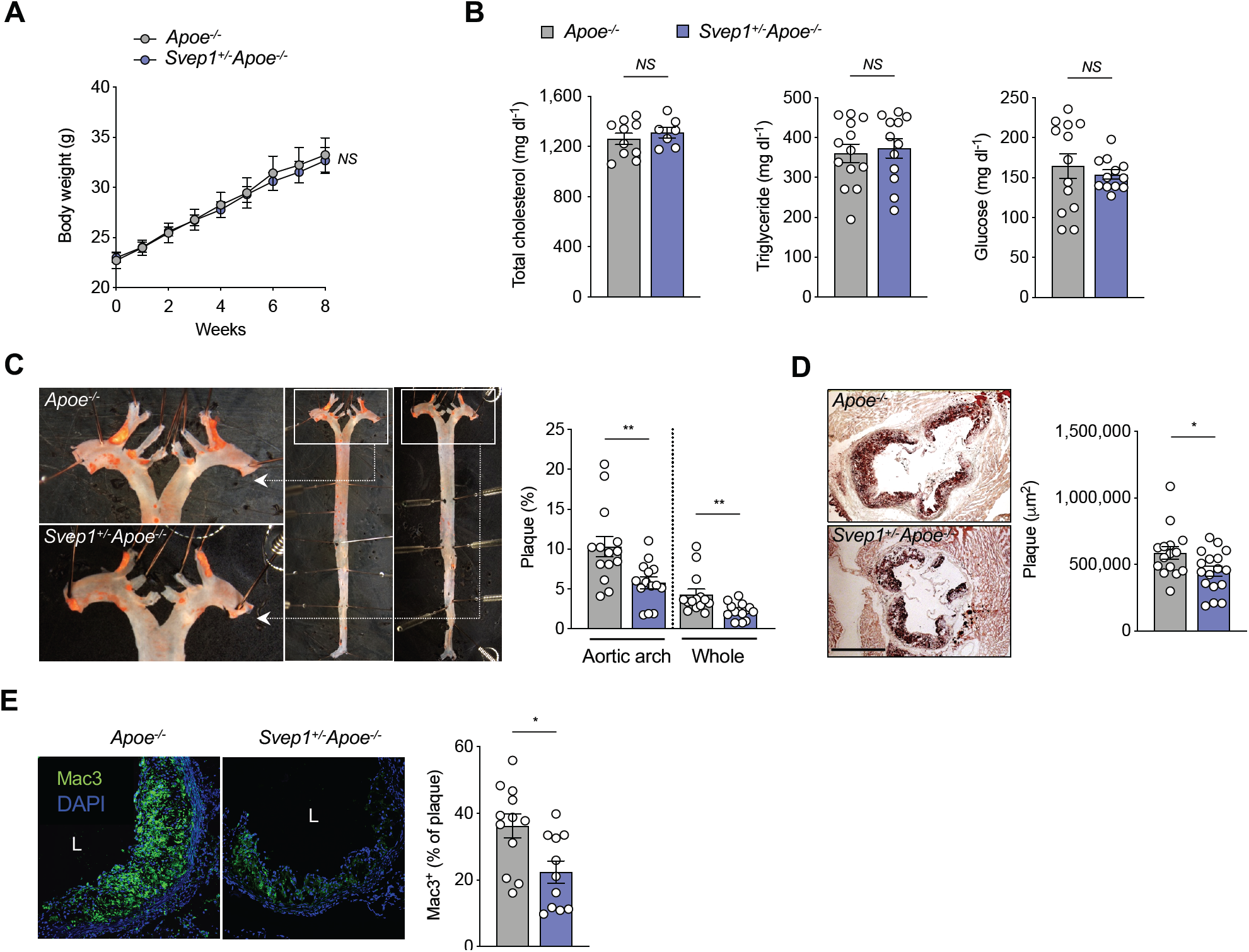
*Svep1* haploinsufficiency abrogates atherosclerosis. (A) Body weight of *Apoe*^*-/-*^ and *Svep1*^*+/-*^*Apoe*^*-/-*^ mice during HFD feeding (*n* = 13-14/group). (B) Plasma total cholesterol (*n* = 7-10), triglycerides, and glucose (*n* = 12-13/group). (C) *En face* Oil Red O-stained aortas. Outlined areas indicate the aortic arch regions magnified in left panels. Quantification of Oil Red O-stained area in each aortic arch and whole artery (*n* = 15-17/group). (D) Oil Red O-stained aortic root cross-sections. Quantification of Oil Red O-stained area (*n* = 15-17/group). Scale bar, 500 µm. (E) Mac3 staining in aortic root sections. Quantification of Mac3 as a percentage of plaque area (*n* = 11-12/group). Scale bar, 200 µm. M, media; L, lumen; P, plaque. One-way ANOVA test (A) or Unpaired nonparametric Mann-Whitney test were used (B through E), and shown as the mean ± SEM. \**P* < 0.05; \*\**P* < 0.01; *NS*, not significant.

We then tested the hypothesis that the atherogenic effects of Svep1 could be attributed to its synthesis by VSMCs using *Svep1*^*SMCΔ/Δ*^ and *Svep1*^*SMC+/+*^ mice, as previously described. As with *Svep1* haploinsufficiency, loss of *Svep1* in VSMCs did not significantly alter body weight, plasma cholesterol, triglycerides and glucose levels (Figure 3A, B) following 8 weeks of HFD feeding. Also consistent with our *Svep1* haploinsufficiency model, *Svep1*^*SMCΔ/Δ*^ mice had decreased plaque burden plaque in the aortic arch, whole aorta, and aortic root (Figure 3C, D), as compared to *Svep1*^*SMC+/+*^ control mice. Additionally, atheromas from *Svep1*^*SMCΔ/Δ*^ mice contained less macrophage staining and necrotic core area, indicators of plaque instability, and unaltered collagen content (Figure S3A-C).

**Figure 3.**
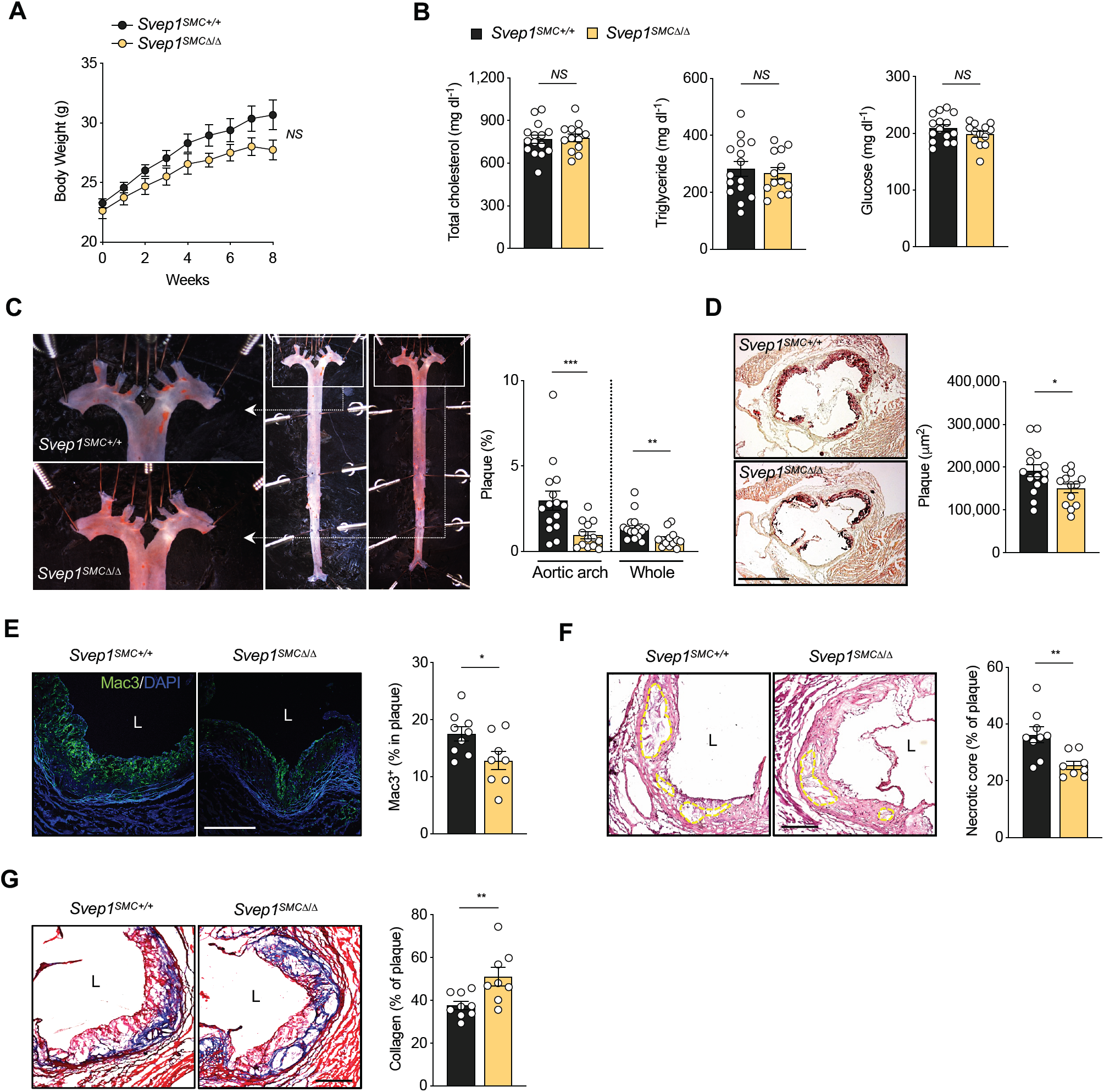
VSMC-specific *Svep1* deficiency reduces atherogenesis and plaque complexity. (A) Body weight of *Svep1*^*SMC+/+*^ and *Svep1*^*SMCΔ/Δ*^ mice during HFD feeding. (B) Total plasma cholesterol, triglycerides, and glucose. (C) *En face* Oil Red O-stained aortas. Outlined areas indicate the aortic arch regions magnified in left panels. Quantification of Oil Red O-stained area in each aortic arch and whole artery. (D) Oil Red O-stained aortic root cross-sections. Quantification of Oil Red O-stained area. Scale bar, 500 µm. *n* = 13-15/group (A through D). (E) Mac3 staining of aortic roots. Quantification of Mac3 as a percentage of plaque area. (F) Necrotic core outlined on H&E-stained sections. (G) Collagen staining using by Masson’s trichrome stain. (E through G) All values were calculated as a percentage of plaque area. Scale bars, 200 µm. *n* = 8-9/group. M, media; L, lumen; P, plaque. One-way ANOVA test (A) or Unpaired nonparametric Mann-Whitney test were used (B through G), and shown as the mean ± SEM. \**P* < 0.05; \*\**P* < 0.01; \*\*\**P* < 0.001; *NS*, not significant.

Given the observations that loss of *Svep1* in VSMCs resulted in a dramatic reduction in plaque size in the setting of 8 weeks HFD feeding, we extended the length of plaque development to investigate the effect of *Svep1* on advanced plaque lesions. After treatment with tamoxifen, *Svep1*^*SMC+/+*^and *Svep1*^*SMCΔ/Δ*^ mice were fed HFD for 16 weeks. Again, no differences were observed in body weight (Figure S3D), plasma cholesterol, and glucose levels (Figure S3E) between groups. Triglycerides were higher in the *Svep1*^*SMCΔ/Δ*^ mice at a level of nominal significance (*P* = 0.046), but this was not observed at other timepoints or in the haploinsufficiency model. Although we did not detect a statistically significant effect of VSMC-specific *Svep1* deletion on atherosclerotic plaque burden (Figure S3F, G), plaques from *Svep1*^*SMCΔ/Δ*^ mice tended to be smaller and were both less complex and more stable than controls. These indicators of an altered plaque phenotype include decreased neointimal macrophage staining (Figure 3E) and necrotic core size (Figure 3F), in addition to greater collagen content (Figure 3G). Taken together, these experimental atherosclerosis data suggest that Svep1 drives atherosclerosis and increases plaque complexity.

### SVEP1 is causally related to cardiometabolic disease in humans

Due to the relationship we discovered between Svep1 depletion and reduced atherosclerosis across our mouse models, we wondered if the human SVEP1 CAD-associated D2072G missense polymorphism was associated with altered SVEP1 levels in humans. While we did not find that this allele (or other alleles in linkage disequilibrium) were associated with mRNA levels in GTEx (Figure S4A), we did find that the 2702G risk variant (SVEP1^CADrv^) was associated with a significant increase in circulating plasma SVEP1 protein levels (*P* = 8 × 10^−14^; Figure 4A) as measured by two independent aptamers (Figure S4B) from participants in the INTERVAL study (Sun et al., 2018), suggesting that increased SVEP1 protein levels were associated with increased risk of CAD. We next wondered if this was true for other genetic variants influencing SVEP1 protein levels. Using published data from the INTERVAL study (Sun et al., 2018), we cataloged *cis*-acting variants that associated with SVEP1 protein levels at a genome-wide (*P* < 5 × 10^−8^) level of statistical significance (Figure 4B). We performed Mendelian Randomization (Burgess et al., 2015) using a subset of these variants in linkage equilibrium (r^2^ < 0.3) and found that increased SVEP1 protein levels were causally related to increased CAD risk (*P* = 7 × 10^−11^; Figure 4C, D). We also asked if SVEP1 protein levels were causally related to increased risk for hypertension and type 2 diabetes due to the prior associations we observed for the SVEP1^CADrv^ allele with these risk factors. Indeed, we found that increased SVEP1 protein levels were causally related to both hypertension (*P =* 2 × 10^−15^; Figure S4D) and type 2 diabetes (*P* = 0.0004; Figure S4E).

**Figure 4.**
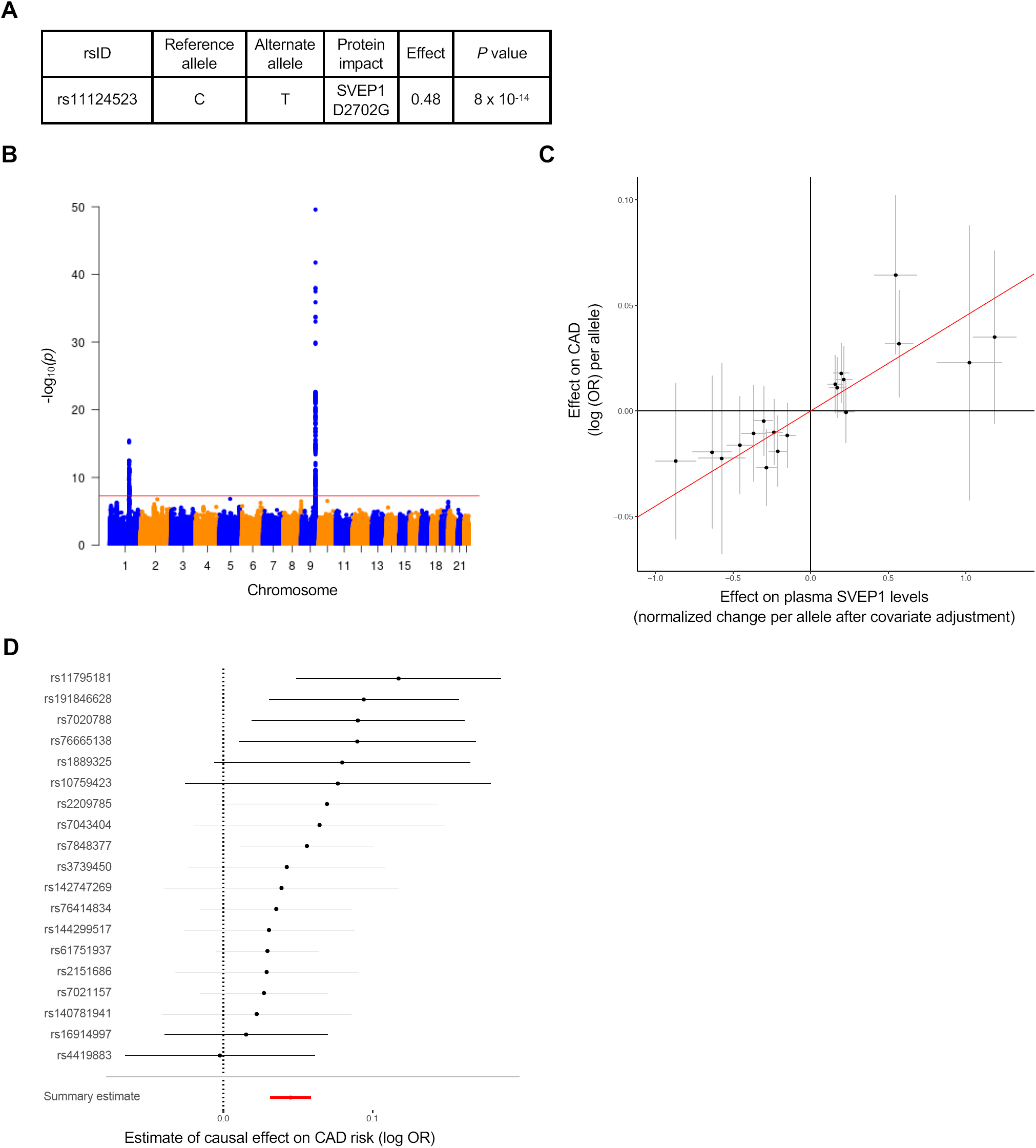
Plasma levels of SVEP1 are causally related to CAD in humans. (A) The effect of the CAD-associated SVEP1 D2702G allele on plasma SVEP1 levels. Effect refers to the change per alternative allele (ending 2702G) in units of normalized protein levels after adjusting for covariates as previously described (Sun et al., 2018). (B) Genome-wide Manhattan plot for variants associated with plasma SVEP1. The –log_10_(p) of the association with SVEP1 levels is plotted for each variant across the genome according to chromosomal position (X-axis). The red line indicates genome-wide significance (*P* < 5 × 10^−8^). The association peak on chromosome 9 overlies the *SVEP1* locus. (C) Estimated effect (with 95% confidence intervals) of each variant included in the Mendelian Randomization analysis on plasma SVEP1 level and CAD risk. The red line indicates the causal effect estimate (*P* = 7 × 10^−11^). (D) The estimated causal effect (with 95% confidence intervals) of each SNP included in the Mendelian Randomization analysis for a one unit increase in SVEP1 level is plotted along with the overall summary estimate from the causal analysis.

To investigate how the human SVEP1^CADrv^ missense polymorphism might impact CAD risk, we generated homozygous mice harboring the human SVEP1^CADrv^ at the orthologous murine position (*Svep1*^*2699G/2699G*^; hereafter referred to as *Svep1*^*G/G*^). These mice were bred with *Apoe*^*-/-*^ mice to generate *Svep1*^*G/G*^*Apoe*^*-/-*^ mice. We were not able to detect differences in body weight, serum total cholesterol, triglycerides, and glucose (Figure S4E-H) between groups after feeding HFD. We also did not appreciate a significant difference between groups in the development of atherosclerotic plaque at either 8 or 16 weeks of HFD feeding (Figure S4I, J). Although our prior human genetic study revealed a significant association with an increased risk of CAD, the effect of the SVEP1^CADrv^ in humans was modest, in which each copy of the G allele was associated with a 14% increased risk of disease. If an effect size in mice is similarly modest, further investigation would require a very large number of animals, presenting both pragmatic and ethical barriers. To circumvent these concerns, subsequent functional interrogation of the SVEP1^CADrv^ was performed in vitro.

### VSMCs express integrin α9β1

To begin characterizing the mechanism by which SVEP1 drives atherosclerosis, we sought to identify receptors and associated cell types that interact with SVEP1 in the extracellular space. Integrin α9β1 is the only protein known to interact with Svep1 and they colocalize *in vivo* (Sato-Nishiuchi et al., 2012). Integrins are transmembrane, heterodimeric receptors that respond to the extracellular environment and influence numerous aspects of atherosclerosis (Misra et al., 2018; Weng et al., 2003). Therefore, we hypothesized that integrin α9β1 (and associated cell-types) may be involved in Svep1-mediated atherosclerosis. ITGA9 is known to exclusively heterodimerize with ITGB1, therefore assessing ITGA9 expression is a reliable proxy for integrin α9β1 expression. Integrin α9β1 expression has been documented in airway epithelium, smooth muscle, skeletal muscle, hepatocytes, and epithelial cells (Chen et al., 2012; Danussi et al., 2011; Gupta and Vlahakis, 2010; Kanayama et al., 2009; Mostovich et al., 2011; Roy et al., 2011; Sato-Nishiuchi et al., 2012; Schreiber et al., 2009), yet arterial tissue expresses the highest *ITGA9* levels of all GTEx tissues (Figure S5A). *In situ* hybridization confirmed that *ITGA9* is broadly expressed in the human aortic wall and LIMA, predominately colocalizing with VSMCs (Figure 5A). Likewise, VSMCs of the murine aorta expressed high levels of *Itgα9* (Figure 5B). Consistent with these data, single cell studies of the murine aorta indicate the VSMCs express *Itgα9* (Figure S5B) (Kalluri et al., 2019). Given the established role of VSMCs in CAD (Bennett et al., 2016), their expression of integrin α9β1, and the local expression patterns of Svep1 in disease, we tested the hypothesis that VSMCs respond to Svep1 in a cell-autonomous manner to promote atherosclerosis.

**Figure 5.**
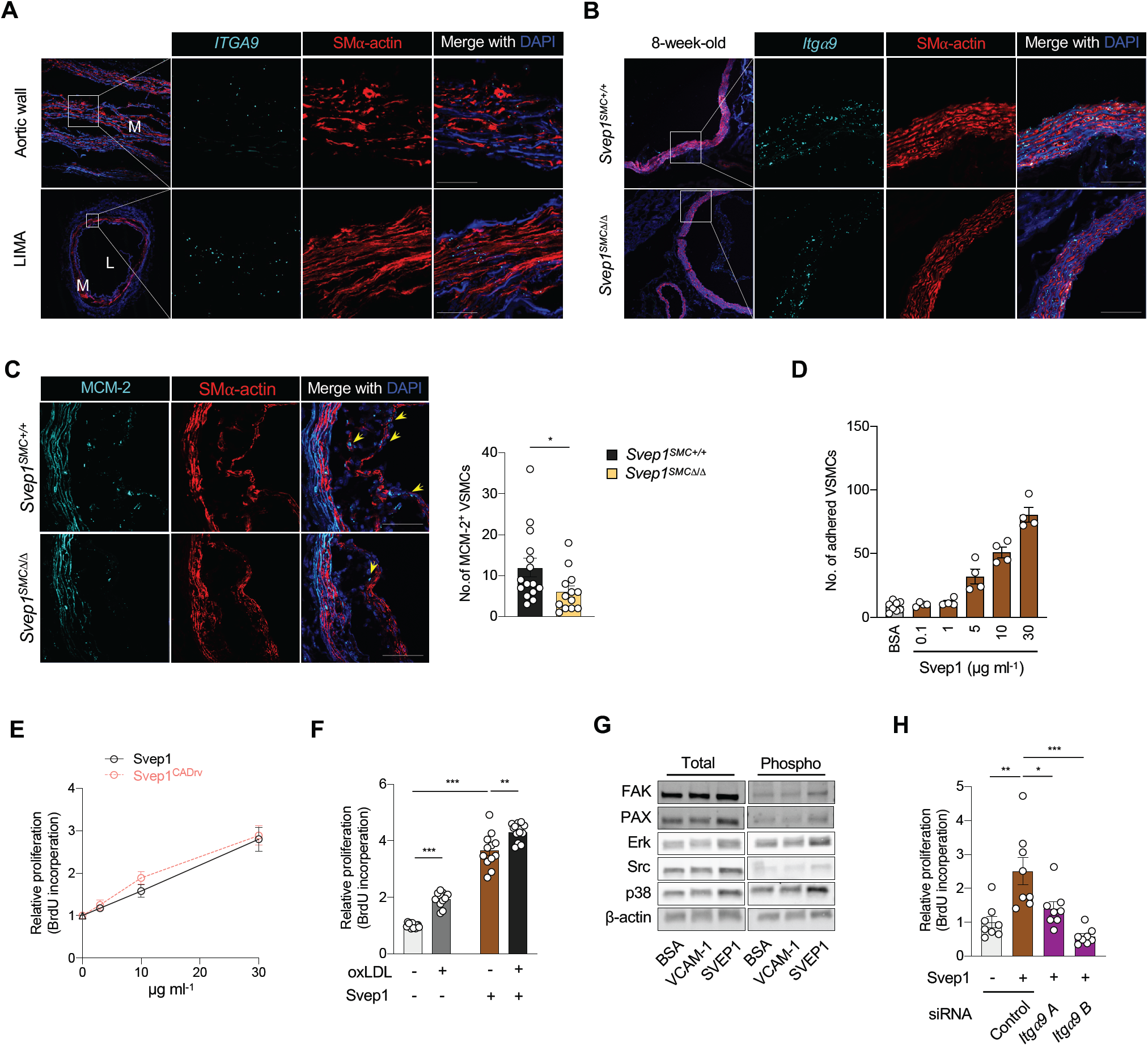
Svep1 induces Itgα9-dependent proliferation in VSMCs. (A) *ITGA9* expression in human aortic wall and LIMA cross-sections from patients using ISH. M, media; L, lumen. (B) Expression of *Itgα9* in the aortic root from 8-week-old *Svep1*^*SMC+/+*^ and *Svep1*^*SMCΔ/Δ*^ mice using ISH. Outlined areas indicate the regions magnified in the next panels. Scale bar, 50 µm. (C) MCM-2 immunofluorescent staining of aortic root regions from *Svep1*^*SMC+/+*^ and *Svep1*^*SMCΔ/Δ*^ mice after 8 weeks of HFD feeding. Yellow arrows indicate MCM-2^+^/SMα-actin^+^ cells within plaque. Quantification of MCM-2^+^/SMα-actin^+^ cells (n = 13-15/group). Scale bars = 50 µm. Tissues in (A-C) were co-stained with the VSMC marker, SMα-actin. (D) Adhesion of VSMCs to increasing concentrations of immobilized Svep1. Adhered cells were counted manually and normalized to wells lacking Svep1. (E) Proliferation of VSMCs in response to increasing concentrations of immobilized Svep1 and Svep1^CADrv^ using a BrdU incorporation assay. (F) *Svep1*^*SMCΔ/Δ*^ VSMCs were incubated in wells precoated with 30 µg ml^-1^ Svep1 protein or BSA (as vehicle control) and treated with or without 50 µg ml^-1^ oxLDL in the culture media for 36 hr. Proliferation was determined by BrdU incorporation. Two-tailed t-test. (G) Immunoblots of integrin signaling kinases and downstream kinases of cells adhered to control, VCAM-1, or Svep1-treated plates. β-actin was used as loading control. (H) VSMCs were transfected with control or *Itga9*-targetted siRNAs and grown on immobilized Svep1 or BSA. Proliferation was determined by BrdU incorporation. \**P* < 0.05; \*\*\**P* < 0.001; \*\*\*\**P* < 0.0001. Two-tailed t-test.

### Svep1 induces proliferation and integrin signaling in VSMCs

The ECM plays a critical role in orchestrating cellular responses to tissue injury, including promoting cell proliferation and differentiation (Bennett et al., 2016; Johnson, 2014). We therefore assessed the proliferation of neointimal *Svep1*^*SMCΔ/Δ*^ and *Svep1*^*SMC +/+*^ VSMCs using immunofluorescent staining of the proliferation marker, mini-chromosome maintenance protein-2 (MCM-2). Among cells expressing smooth muscle actin, fewer stained positive for MCM-2 in *Svep1*^*SMCΔ/Δ*^ mice as compared to *Svep1*^*SMC +/+*^ controls after HFD feeding for 8 weeks (Figure 5C), suggesting Svep1 induces VSMC proliferation.

To further explore the effects of Svep1 on VSMCs, we generated and purified recombinant Svep1 and its orthologous CAD risk variant (Svep1^CADrv^) using a mammalian expression system. We tested the response of primary VSMCs to Svep1 that was immobilized on culture plates, reflecting an overexpression-like assay while maintaining its physiologic context as an extracellular matrix protein (in contrast to genetic overexpression). VSMCs adhere to Svep1 in a dose dependent manner (Figure 5D). Exposure to both Svep1 variants induces dose-dependent VSMC proliferation, based on BrdU incorporation (Figure 5E). As a point of reference, we used oxLDL, a proliferative stimulus relevant to atherosclerosis, in addition to Svep1 to test VSMC proliferation. Strikingly, Svep1 was able to induce more VSMC proliferation than oxLDL. Exposure to a combination of oxLDL and Svep1, as exists within the atheromatous environment, caused the greatest amount of VSMC proliferation (Figure 5F). Murine macrophages exposed to Svep1 do not proliferate in the absence or presence of oxLDL (Figure S5C) suggesting that Svep1 is not a proliferative stimulus for all cell types.

Integrin α9β1 is expressed by VSMCs, binds to Svep1, and drives proliferation in some cell types (Schreiber et al., 2009). Therefore, to begin to interrogate the molecular mechanisms by which Svep1 influences VSMCs, we tested whether Svep1 exposure was able to induce integrin signaling in VSMCs. We tested this by seeding cells to wells coated with bovine serum albumin (as an inert protein control), VCAM-1 (a low affinity integrin α9β1 ligand), or Svep1 (a high affinity integrin α9β1 ligand). We found that cells adherent to Svep1 had increased phosphorylation of canonical integrin signaling kinases, such as focal adhesion kinase (FAK), Paxillin (Pax), and Src, as well as downstream MAPK kinases, ERK and p38 (Figure 5G), relative to an inert protein control. Svep1^CADrv^ had similar effects as Svep1 on integrin signaling in VSMCs (Figure S5D). We then tested if Svep1-induced proliferation was dependent on integrin α9β1. Since Itgα9 exclusively heterodimerizes with Itgβ1, we used siRNA knockdown of *Itgα9* to disrupt integrin α9β1. The proliferative effect of Svep1 was completely inhibited by knockdown of *Itgα9* using two different siRNA constructs (Figure 5H), suggesting that integrin α9β1 is necessary for Svep1-induced VSMC proliferation.

### Svep1 regulates key VSMC differentiation pathways

We next sought to characterize the response of primary VSMCs to the wildtype Svep1 and Svep1^CADrv^ proteins using an unbiased methodology. Cells were collected after 20 hours of growth on the indicated substrate and transcriptomic analysis was performed using RNA-sequencing. Pathway and gene ontology analysis was used to determine the shared and unique transcriptional response to the Svep1 variants. Consistent with previous findings, a number of cell adhesion and proliferation-related pathways and terms were enriched in the shared transcripts of cells exposed to either Svep1 variant. These include ECM-receptor interaction, focal adhesion, integrin-mediated signaling, positive regulation of cell proliferation, and various additional proliferative and mitogenic pathways (Figure 6A, B, Supplemental Table 1). A striking number of differentiation and development-related pathways and terms were also enriched in cells exposed to the Svep1 variants. These include angiogenesis, cell differentiation, and wound healing, among many others (Figure 6A, B, Supplemental Table 1).

**Figure 6.**
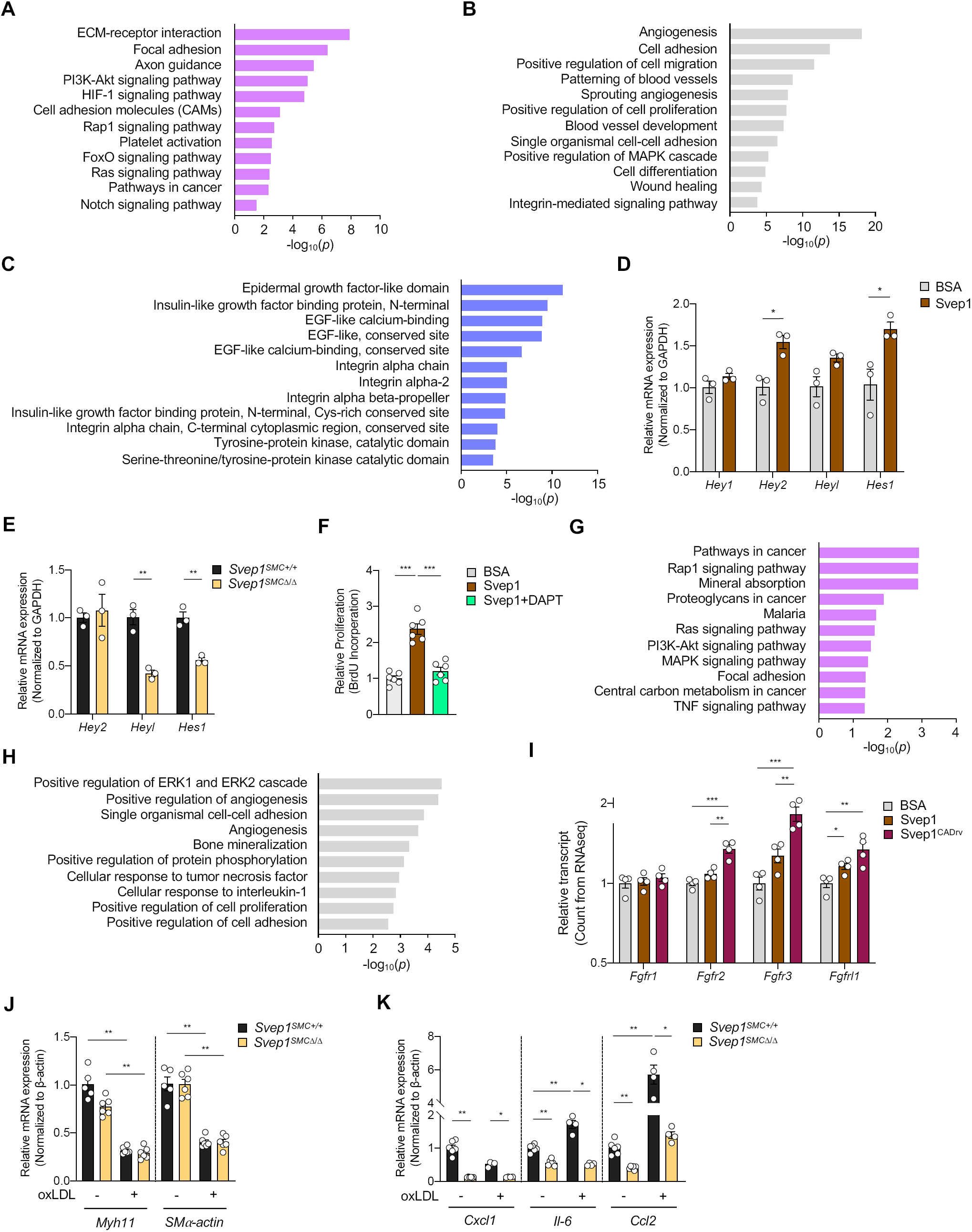
Svep1 modulates key VSMC-developmental pathways. (A-C) Common transcriptional response of VSMCs to Svep1 and Svep1^CADrv^ proteins. Dysregulated (A) KEGG pathways, (B) GO term molecular functions, and (C) InterPro domains. Top 5 dysregulated categories plus additional, select categories are included. Full results are available in Supplemental Table 1. Bars represent -log_10_ of *P* values. (D) Transcription of canonical Notch target genes after 4 hours of adhesion to Svep1, relative to BSA. Two-tailed t-test. (E) Basal transcription of Notch target genes in *Svep1*^*SMC+/+*^ and *Svep1*^*SMCΔ/Δ*^ VSMCs. Two-tailed t-test. (F) Proliferation of VSMCs in response to immobilized Svep1. Cells were treated with DMSO (carrier) or 25 µM DAPT. Proliferation was determined by BrdU incorporation. Two-tailed t-test. (G-H) Differential transcriptional response of VSMCs to Svep1 and Svep1^CADrv^ proteins. Dysregulated (A) KEGG pathways, (B) GO term molecular functions. Top 5 dysregulated categories plus additional, select categories are included. Full results are available in Supplemental Table 1. Bars represent -log_10_ of *P* values. (I) Bar graph of *Fgfr* transcript counts from RNAseq. Each transcript is normalized to the BSA control group. Two-tailed t-test. (J, K) qPCR of (J) VSMC markers, and (K) inflammatory markers of VSMC cultured with or without 50 µg ml^-1^ oxLDL for 24 hr. \**P* < 0.05; \*\**P* < 0.01; \*\*\**P* < 0.001.

Svep1 contains numerous different and repeating domains that are known to play critical developmental roles and may therefore be governing the effects of Svep1 on VSMCs. Further, although *Svep1*^*-/-*^ and *Itgα9*^*-/-*^ mice have similar phenotypes of edema and lymphatic defects (Karpanen et al., 2017; Morooka et al., 2017), the phenotype of *Svep1*^*-/-*^ mice is markedly more severe (death by E18.5 vs P12 (Huang et al., 2000)), suggesting Itgα9 may have partial redundancy with an additional receptor(s) for Svep1. To search for evidence of additional domain interactions, we cross-referenced the transcriptional profile of VSMCs to the Svep1 variants with InterPro (Mitchell et al., 2019), a database of protein domains. In addition to integrin-related domains, transcripts that code for EGF-like domain-containing proteins were highly differentially expressed in cells exposed to Svep1 (Figure 6C). Repeat EGF-like domains often interact, as occurs in Notch signaling, suggesting Svep1’s repeat EGF-like domains may be playing an important, but as of yet undescribed role in the biological function of Svep1 (Mitchell et al., 2019). Indeed, transcripts related to Notch signaling were dysregulated in cells exposed to Svep1 (Figure 6A).

As an orthogonal approach to interrogating SVEP1’s mechanisms and potential binding partners, we sought to identify homologues in distantly related species. The Drosophila protein, uninflatable, is a potential orthologue of SVEP1 (Sonnhammer and Ostlund, 2015) and contains a region defined by three ephrin-receptor like domains, followed by tandem EGF-repeats and a Laminin-G domain (Marchler-Bauer et al., 2011), mirroring a region of SVEP1 that contains a highly similar sequence of domains. Inhibition of uninflatable in Drosophila larvae results in defective tracheal development, analogous to the vascular defects observed in zebrafish Svep1 mutants (Ghabrial and Krasnow, 2006; Zhang and Ward, 2009). Uninflatable has been shown to bind and modulate Notch signaling in Drosophila (Loubery et al., 2014; Xie et al., 2012; Zhang and Ward, 2009). These findings, in addition to the RNAseq analysis, led us to hypothesize that Svep1 may also modulate Notch signaling.

VSMCs express multiple Notch receptors (Davis-Knowlton et al., 2019), thus, we tested the impact of Svep1 on Notch signaling in VSMCs. This was assessed by seeding VSMCs on tissue culture plates treated with Svep1 or BSA (as an inert control protein) for 4 hours, since Notch signaling is highly temporally regulated (Schweisguth, 2004). Cells grown on Svep1 had significantly increased expression of canonical Notch targets *Hey2* and *Hes1* even without overexpression of a Notch receptor (Figure 6D). Conversely, primary VSCMs collected from *Svep1*^*SMCΔ/Δ*^ mice have decreased transcription of Notch target genes (Figure 6E), supporting the regulation of Notch signaling by Svep1. Svep1-induced proliferation was also completely abrogated upon Notch inhibition by the γ-secretase inhibitor, DAPT (Figure 6F). Cell proliferation in response to 10% fetal bovine serum was not significantly inhibited by DAPT (data not shown), demonstrating that Notch signaling is necessary for Svep1-induced proliferation. It is possible that Notch and integrin receptors may cooperatively regulate the effects of SVEP1, similar to that reported on non-canonical ECM Notch regulators MAGP2 and EGFL7 (Deford et al., 2016).

### VSMCs differentially respond to Svep1^CADrv^ compared to Svep1

Our experimental atherosclerosis models and Mendelian Randomization analysis indicate that both SVEP1 variants are atherogenic, with Svep1^CADrv^ having the greater atherogenicity of the two. We therefore interrogated the differential transcriptional responses of VSMCs to the Svep1 variants. This analysis also revealed that a large number of proliferation-related pathways were disproportionately regulated by the variants (Figure 6G, H, Supplementary Table 1). Further exploration revealed that the fibroblast growth factor (FGF) receptor family was differentially expressed between the variants. The FGFR family is also sub-categorized within several of the most differentially regulated pathways and terms. FGF signaling is proatherogenic in VSMCs (Chen et al., 2016), so we assessed the effect of each variant on the direction and magnitude of transcription of each FGF receptor expressed by VSMCs. Consistent with their relative atherogenicities, SVEP1 increases expression of FGF receptors but exposure to Svep1^CADrv^ resulted in significantly higher expression of FGF receptors (Figure 6I). These data suggest that increased FGF signaling may contribute to the increased CAD risk associated with SVEP1^CADrv^.

Given the fundamental role of integrin, Notch, and FGFR signaling in regulating VSMC phenotype, we assessed the effects of Svep1 in response to oxLDL, an inflammatory stimulus relevant to atherosclerosis. Upon oxLDL stimulation, both *Svep1*^*SMC+/+*^ and *Svep1*^*SMCΔ/Δ*^ VSMCs decreased the expression of contractile markers *Myh11* and *SMα-actin* (Figure 6J), while increasing expression of the inflammatory markers *Il-6* and *Ccl2* (Figure 6K), confirming an inflammatory response to oxLDL. *Cxcl1, Il-6*, and *Ccl2* expression was lower in *Svep1*^*SMCΔ/Δ*^ VSMCs than *Svep1*^*SMC+/+*^ controls, suggesting that Svep1 may be a pro-inflammatory stimulus VSMCs under atherosclerotic conditions.

### Svep1 promotes inflammation in atherosclerosis

To investigate how the loss of *Svep1* influences pathways involved in the development of atherosclerosis at the tissue level, we performed RNA-seq analyses on mRNA extracted from aortic arches of *Svep1*^*SMC+/+*^and *Svep1*^*SMCΔ/Δ*^ mice after 8 weeks of HFD. Loss of *Svep1* in VSMCs altered a number of inflammatory pathways upon induction of atherosclerosis. These include cytokine-cytokine receptor interaction, chemokine signaling, and NF-kappa B signaling pathways (Figure 7A, B). Notably, both cell adhesion molecules (CAMs) and ECM-receptor interaction were also dysregulated in the atherosclerotic aortic arches from *Svep1*^*SMCΔ/Δ*^ (Figure 7A, B and Supplementary Table 2). Quantitative PCR using cDNA from the aortic arches of the same mice was used to validate the RNA-seq results. Specifically, *Ccl2* (C-C motif chemokine ligand 2), *Spp1* (secreted phosphoprotein 1, also known as osteopontin), and *Cxcl5* (C-X-C motif chemokine ligand 5) were significantly decreased in *Svep1*^*SMCΔ/Δ*^ mice, as compared to *Svep1*^*SMC+/+*^ mice (Figure S6A). Despite these differences, we did not find a significant alteration in circulating inflammatory mediators in these mice, suggesting Svep1 influences local tissue inflammation but not systemic inflammation (Figure S6B). These data are also consistent with our observations that *Svep1* depletion decreases neointimal macrophage staining in atherosclerotic plaque.

**Figure 7.**
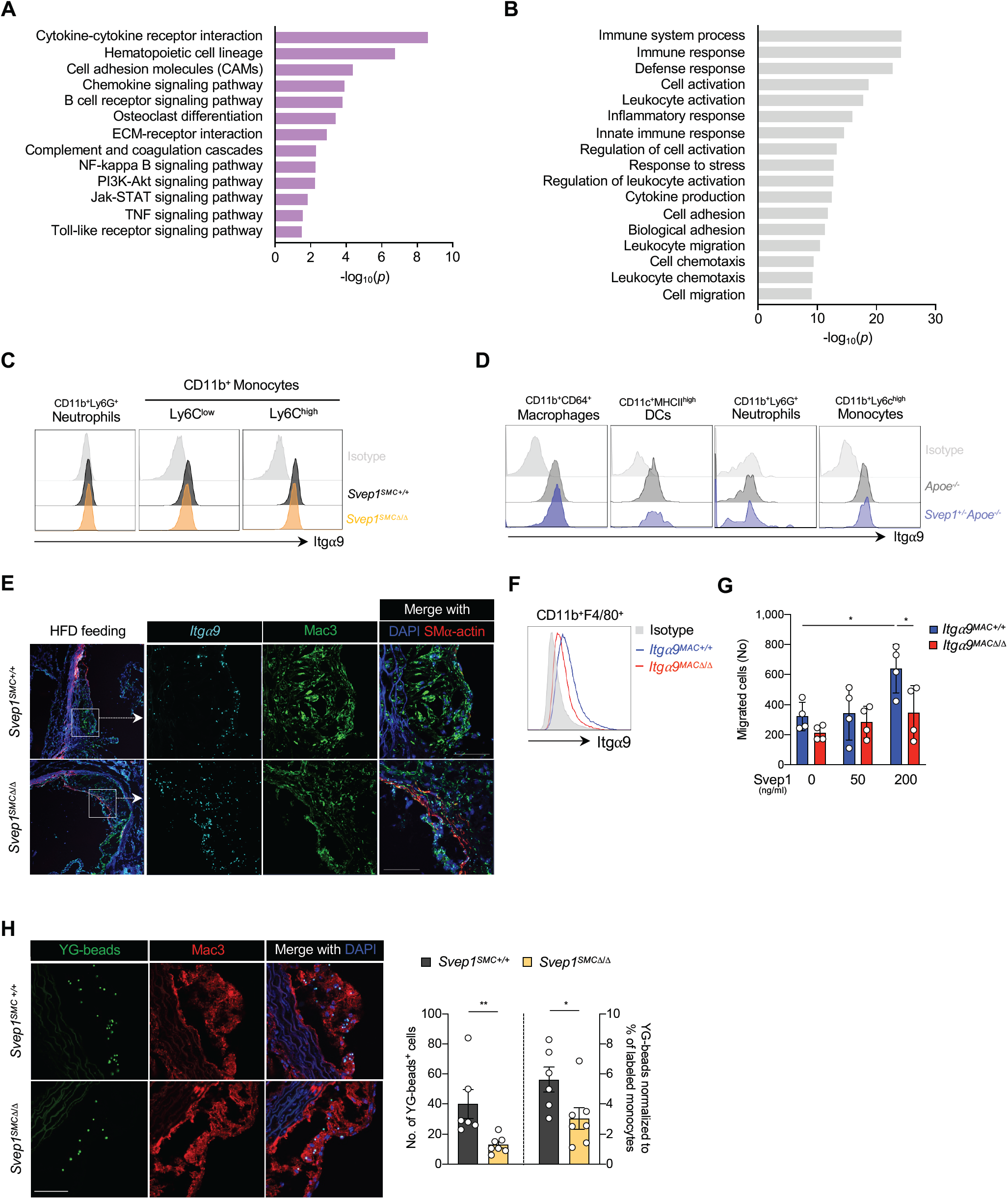
Svep1 promotes inflammation in atherosclerosis. (A-B) Differential transcriptional profile of atherosclerotic aortic arches from *Svep1*^*SMC+/+*^ and *Svep1*^*SMCΔ/Δ*^ mice. Dysregulated (A) KEGG pathways (B) GO term molecular functions. Top 5 dysregulated categories plus additional, select categories are included. Full results are available in Supplemental Table 2. Bars represent -log_10_ of *P* values. (C) Histogram for *Itgα9β1* expression in mouse blood neutrophils (CD11b^+^Ly6G^+^), Ly6C^low^ (CD11b^+^Ly6C^low^), and Ly6C^high^ (CD11b^+^Ly6C^high^) monocytes from *Svep1*^*SMC+/+*^ and *Svep1*^*SMCΔ/Δ*^ mice after 8 weeks of HFD. (D) Histogram of *Itgα9β1* expression in the subpopulations of aortic leukocytes. Macrophages (CD64^+^CD11b^+^), DCs (CD11c^+^MHCII^high^), neutrophils (CD11b^+^Ly6G^+^), and Ly6C^high^ (CD11b^+^Ly6C^high^) monocytes from *Apoe*^*-/-*^ and *Svep1*^*+/-*^*Apoe*^*-/-*^ mice after 8 weeks of HFD. (E) Expression of *Itga9* in the aortic roots from *Svep1*^*SMC+/+*^ and *Svep1*^*SMCΔ/Δ*^ mice using ISH after 8 weeks of HFD. Tissues were co-stained for Mac3 and SMα-actin. Scale bars, 50 µm. (F) Expression of Itgα9 in BMDM from *Itgα9*^*MAC+/+*^ and *Itgα9*^*MACΔ/Δ*^ mice. (G) Migratory response of thioglycolate-elicited macrophages from *Itgα9*^*MAC+/+*^ and *Itgα9*^*MACΔ/Δ*^ were determined using a chemotaxis chamber incubated with 0, 50, and 200 ng ml^-1^ of Svep1 protein. Migrated cells were counted by an automated microscope and expressed as cells per field of view. (H) *In vivo* monocyte recruitment assay. YG-bead uptake within plaque lesion in the aortic root regions from *Svep1*^*SMC+/+*^ and *Svep1*^*SMCΔ/Δ*^ mice. Quantification of YG-bead uptake showing the total number of YG-beads per section (left Y axis), and the number of YG-beads normalized to the percentage of labeled Ly6C^low^ monocytes (right Y axis). *n* = 6-7/group. Scale bar, 50 µm. \**P* < 0.05; \*\**P* < 0.01.

Integrins play a critical role in the immune response, we therefore asked whether immune cells may also express integrin α9β1 and interact with SVEP1 in atherosclerosis. In human peripheral blood cells, moderate integrin α9β1 expression was detected by neutrophils and low expression was detected by CD14^low^CD16^+^ non-classical, CD14^high^CD16^+^ intermediate, and CD14^+^CD16^-^ classical monocytes (Figure S7A) as previously reported (Shang et al., 1999). Given that monocytes significantly alter their expression profiles upon tissue entry and differentiation into macrophages (Chistiakov et al., 2015), we sought to test if macrophages in atherosclerotic plaque express *ITGA9*. Indeed, *ITGA9* expression was detected in CD68^+^ macrophages within human atherosclerotic plaque by *in situ* hybridization (Figure S7B).

We then sought to further assess the expression of integrin α9β1 expression in circulating murine leukocyte subsets. High expression of integrin α9β1 was detected in both Ly6C^hi^ and Ly6C^low^ monocytes and we could detect low levels in neutrophils (Figure 7C). These expression patterns were unaltered in heterozygous *Svep1* deficiency (Figure S7C) and we did not observe an induction of integrin α9β1 expression upon oxLDL treatment in any cell type tested (Figure S7D, E). Considering the finding that integrin α9β1 is expressed by monocyte subsets in peripheral mouse blood, we further analyzed its expression in myeloid cells from the aortas of *Apoe*^*-/-*^ and *Svep1*^*+/-*^*Apoe*^*-/-*^ mice following 8 weeks of HFD feeding. We discovered that integrin α9β1 was expressed in both macrophages and Ly6C^hi^ monocytes of these mice (Figure 7D), consistent with human expression data. We similarly detected robust expression of *Itgα9* by neointimal macrophages using *in situ* hybridization (Figure 7E).

Since integrin α9β1 is expressed on monocytes/macrophages, we sought to better understand whether Svep1 could be directly interacting with integrin α9β1 on these cells. To test this, we generated mice with myeloid cell lineage-specific knockout of *Itg*α*9* using *LysM-Cre* (*Itgα9*^*flox/flox*^*LysM-Cre*, hereafter referred to as *Itga9*^*MACΔ/Δ*^). *Itga9*^*+/+*^*LysM-Cre* mice, referred to as *Itga9*^*MAC+/+*^, served as controls. First, we confirmed that bone marrow derived macrophages from *Itga9*^*MACΔ/Δ*^ animals had a significant reduction in the amount of integrin α9β1 that was present on the cell surface (Figure 7F). We then tested the ability of peritoneal macrophages from these animals to migrate in response to Svep1 using a trans-well migration assay. Svep1 exposure induced a dose-dependent trans-well migration of macrophages from *Itga9*^*MAC+/+*^ control animals but not from *Itga9*^*MACΔ/Δ*^ mice (Figure 7G). This suggests that Svep1 and integrin α9β1 may directly interact to augment myeloid cell homing or migration. Consistent with this, THP-1 cells, a human monocytic cell line, adhered to Svep1 in a dose-dependent manner (Figure S7F, G). Integrin signaling was also activated in THP-1 cells upon exposure to Svep1 or Svep1^CADrv^ and no differences were observed between the variants (Figure S7G).

To test if Svep1 had similar effects on leukocytes *in vivo*, we performed an *in vivo* monocyte recruitment assay in *Svep1*^*SMC+/+*^ and *Svep1*^*SMCΔ/Δ*^ mice. After 8 weeks of HFD feeding, we injected yellow/green (YG) latex beads intravenously in order to label circulating Ly6C^low^ monocytes. Flow cytometry was performed three days after intravenous bead injection (to confirm labeling) and the aortic tissues were isolated for histology on the fourth day following bead injection (to assess recruitment). We confirmed that YG beads were preferentially labeled on Ly6C^low^ monocytes and not on Ly6C^high^ monocytes, indicating efficient bead labeling of circulating monocytes (Figure S7H). We did not observe a difference between groups in the efficiency of bead labeling for monocyte subsets (Figure S7I). Next, we quantified the number of labeled monocytes recruited into atherosclerotic plaques of aortic roots using fluorescent microscopy. *Svep1*^*SMCΔ/Δ*^ mice had significantly fewer YG beads per atheroma, with or without normalization to the percentage of labeled monocytes, relative to *Svep1*^*SMC+/+*^ mice (Figure 7H).

Taken together, these data support Svep1’s role in promoting inflammation in atherosclerosis, either indirectly by promoting an inflammatory VSMC phenotype, directly by interacting with integrin α9β1 on circulating or tissue leukocytes, or a combination of these processes.

## Discussion

Human genomic studies hold great promise in identifying therapeutic targets for disease (Young and Stitziel, 2019), but a significant limitation in translating their findings is the identification of specific causal genes that underlie the observed statistical associations. In a previous study, we identified a low-frequency polymorphism in *SVEP1* that robustly associated with coronary artery disease risk in humans (Stitziel et al., 2016), but it was not clear if *SVEP1* was the causal gene in the locus. Here, we present the first report that SVEP1 is causal in coronary artery disease using experimental mouse models and Mendelian Randomization.

Atherosclerosis is a complex, multifactorial disease process with numerous cell types playing a role in its pathogenesis. This presents an arduous challenge when validating genomic risk loci and testing their mechanisms. The SVEP1^CADrv^ does not associate with changes in plasma lipid levels (Stitziel et al., 2016), prompting us to explore how SVEP1 might influence other aspects of disease pathogenesis. We used human and mouse expression data at the cell and tissue level to develop mechanistic hypotheses, which we then tested using *in vivo* and *in vitro* approaches. Specifically, high basal arterial expression of both *SVEP1* and *ITGA9*, and increased SVEP1 expression under pathological conditions, led us to hypothesize that these proteins may influence local disease processes. Upon exposure to various pathologic stimuli, VSMCs can undergo a “phenotype shift”, in which they lose their quiescent, contractile properties and become migratory, proliferative, inflammatory, and synthetic (Basatemur et al., 2019; Bennett et al., 2016). VSMCs gain properties of matrix-synthesizing fibroblasts during atherosclerosis (Wirka et al., 2019), making VSMCs our primary candidates for the source of SVEP1 within atherosclerotic plaque. Our results provide strong evidence that atherogenic SVEP1 is indeed synthesized by VSMC-derived cells within the atherosclerotic plaque.

We then used expression of *ITGA9* to identify disease-relevant cell types that may respond to SVEP1. This led to the hypothesis that SVEP1 may be interacting with VSMCs by an autocrine mechanism or monocytes by a paracrine mechanism to promote atherosclerosis. VSMCs play a particularly complex and intriguing role in atherosclerosis and warrant further discussion. Recent lineage tracing studies have challenged the notion that VSMCs play a protective role in atherosclerosis (Bennett et al., 2016) by demonstrating that a large, heterogenous population of cells within plaque are derived from VSMCs (Basatemur et al., 2019; Bennett et al., 2016; Shankman et al., 2015). Furthermore, numerous CAD risk loci have now been linked to VSMCs (Liu et al., 2018). This study demonstrates that Svep1 profoundly influences the behavior of VSMCs by regulating a number of pathways with vital roles in VSMC biology. These pathways include integrin, Notch, and FGFR signaling, each of which has been shown to contribute to atherosclerosis (Boucher et al., 2012; Chen et al., 2016; Fukuda et al., 2012; Misra et al., 2018). Recent studies have provided novel insights into the regulation of VSMC phenotype in atherosclerosis by various transcription factors (Cherepanova et al., 2016; Shankman et al., 2015; Wirka et al., 2019). The ECM also plays a fundamental role in regulating VSMC phenotype and is amenable to pharmacologic intervention. Current strategies for the treatment and prevention of CAD consist of lowering risk factors, such as plasma lipids, yet substantial residual risk remains despite effective treatment. Intervening on VSMCs may be a powerful complimentary approach to these traditional therapies.

In addition to its association with CAD, our Mendelian randomization results suggest that circulating SVEP1 causally underlies risk of hypertension and type 2 diabetes. Although the source of SVEP1 in human plasma is unknown, other ECM proteins have been detected in the circulation of patients with atherosclerosis, suggesting that plasma levels of these proteins may reflect tissue levels and atherosclerotic remodeling (Langley et al., 2017; Sundstrom and Vasan, 2006). The mechanisms by which the genetic variants used in the Mendelian randomization affect plasma SVEP1 levels is unclear. Two reasonable hypotheses include modification of protein secretion or degradation, however further studies will be required to determine these mechanisms. Regardless, the power of the two sample Mendelian randomization framework is that these alleles are allocated randomly at birth and are associated with SVEP1 levels in the absence of disease, suggesting that the presence of disease is not driving altered SVEP1 levels, but rather that altered SVEP1 levels are causally related to disease. This further suggests that circulating SVEP1 levels have the potential to be useful as a predictive biomarker.

Additional human genetic data also supports a broader role of SVEP1 in cardiometabolic disease. The alpha subunit of integrin α9β1, which binds to SVEP1 (Sato-Nishiuchi et al., 2012) with an affinity that far exceeds its other known ligands (Andrews et al., 2009; Hakkinen et al., 2000; Nishimichi et al., 2009; Smith et al., 1996), is also associated with blood pressure in multiple studies (Levy et al., 2009; Takeuchi et al., 2010). Overexpression of disintegrin and metalloproteinase with thrombospondin motifs-7 (ADAMTS-7), another CAD risk locus, in primary rat VSMCs alters the molecular mass of SVEP1 (Kessler T, 2015). The overlapping disease associations and molecular interactions between these three risk loci converge on SVEP1 and point to a regulatory circuit with a prominent, yet unexplored role in cardiometabolic disease. Further studies will be required to validate their interactions and mechanisms *in vivo*, and to explore the potential of targeting this pathway for the treatment of cardiometabolic disease.

Our complementary mouse models demonstrate that *Svep1* haploinsufficiency and VSMC-specific *Svep1* deficiency significantly abrogate the development of atherosclerosis. Each intervention was well tolerated by mice, as we did not observe any adverse response to Svep1 depletion. Similarly, our Mendelian Randomization analyses suggest there may be a therapeutic window to safely target SVEP1 levels. These findings suggest that targeting SVEP1 or selectively modulating its interactions may be a viable strategy for the treatment and prevention of coronary artery disease.

## Acknowledgements

This work was supported in part by grants from the National Institutes of Health (NIH) T32GM007200 and T32HL134635 to JSE and NIH T32HL007081 to EPY, along with NIH grants R01HL131961, UM1HG008853, UL1TR002345, an investigator-initiated research grant from Regeneron Pharmaceuticals, a career award from the National Lipid Association, and by the Foundation for Barnes-Jewish Hospital to NOS. We thank the Genome Technology Access Center in the Department of Genetics at Washington University School of Medicine for help with genomic analysis. The Center is partially supported by NCI Cancer Center Support Grant #P30 CA91842 to the Siteman Cancer Center and by ICTS/CTSA Grant# UL1TR002345 from the National Center for Research Resources (NCRR), a component of the National Institutes of Health, and NIH Roadmap for Medical Research. We thank the Bursky Center for Human Immunology and Immunotherapy Programs at Washington University, Immunomonitoring Laboratory for help with analysis of mouse plasma. This publication is solely the responsibility of the authors and does not necessarily represent the official view of NCRR or NIH. The Svep1 mouse strain used for this research was created from ES cell clone HEPD0747_6_B06, generated by the European Conditional Mouse Mutagenesis Program which was then made into mice and provided to the KOMP Repository (www.komp.org) by the Jackson Laboratory as part of the KOMP2 Project. The Genotype-Tissue Expression (GTEx) Project was supported by the Common Fund of the Office of the Director of the National Institutes of Health, and by NCI, NHGRI, NHLBI, NIDA, NIMH, and NINDS. Data used for the analyses described in this manuscript were obtained from the GTEx Portal on 04/30/2020. We thank all the members of the Stitziel Lab for helpful discussion.

## Author information

NOS conceived of the study. IHJ performed animal experiments. JSE and IHJ performed *in vitro* experiments. IHJ, JSE, AA, and NOS designed and interpreted the experiments. AA and KS generated critical reagents. EPY and CJK performed Mendelian Randomization analyses. PK provided and assisted with human specimens. KJL, BR, and RPM provided expertise in animal models and data interpretation. IHJ, JSE, and NOS wrote the manuscript. All authors reviewed and provided critical editing of the manuscript.

## Methods

### Human tissue collection

Prior to coronary artery bypass grafting surgery (CABG) for the treatment of symptomatic coronary artery disease, we consented five patients for tissue and peripheral blood collection at the time of their planned CABG to be performed at Barnes Jewish Hospital. The surgical plan for all patients included using the left internal mammary artery (LIMA) as an arterial graft to the left anterior descending (LAD) coronary artery and at least one venous graft to a different coronary artery. During the CABG, we collected the distal end of the LIMA which was trimmed in order to accommodate the length needed to reach the LAD. We also collected the aortic wall punch biopsy that was used to provide a proximal anastomotic site for the venous conduit. Tissues were immediately placed in phosphate buffered saline (PBS) on ice and brought to the laboratory where they were frozen at -80°C prior to in situ hybridization described below. During the CABG, we also collected 5-7 ml of peripheral blood in a tube containing the anticoagulant K_3_ EDTA (#6457, BD Biosciences) which was used for flow cytometry as described below. All research participants provided written informed consent and the study was approved by the Washington University School of Medicine Human Research Protection Office and Institutional Review Board.

### Mice

All animal studies were approved by the Animal Studies Committee and the Institutional Animal Care and Use Committee of the Washington University School of Medicine. *Svep1*^*+/-*^ mice were made by KOMP (knockout mouse project), and these mice were then crossed with mice expressing the flippase FLP recombinase under the control of the promoter of the human actin beta gene (hATCB) to generate *Svep1*^*flox/flox*^ (*Svep1*^*Δ/Δ*^) mice. CRISPR/Cas9 genome editing technology was used in collaboration with the Washington University School of Medicine Genome Engineering and Transgenic Micro-Injection Cores to generate *Svep1*^*G/G*^ mice on a C57BL/6 background harboring the *SVEP1* mutation at the homologous murine position (p.D2699G). *Svep1*^*+/-*^ and *Svep1*^*G/G*^ mice were crossed with *Apoe*^*-/-*^ mice (#002052, Jackson Laboratory) to get *Svep1*^*+/-*^*Apoe*^*-/-*^ and *Svep1*^*G/+*^*Apoe*^*-/-*^ mice, which we maintained as breeders to generate experimental and control mice. We crossed *Svep1*^*Δ/Δ*^ mice with *Myh11-CreER*^*T2*^ (#019079, Jackson Laboratory) mice to generate *Svep1*^*Δ/+*^*Myh11-CreER*^*T2*^ mice. *Svep1*^*Δ/+*^*Myh11-CreER*^*T2*^ males were then crossed with *Svep1*^*Δ/+*^ females to generate experimental *Svep1*^*Δ/Δ*^*Myh11-CreER*^*T2*^ and control *Svep1*^*+/+*^*Myh11-CreER*^*T2*^ male littermate mice. Finally, *Svep1*^*Δ/Δ*^*Myh11-CreER*^*T2*^ males were crossed with *Apoe*^*-/-*^ females. We maintained *Svep1*^*Δ/+*^*Myh11-CreER*^*T2*^*Apoe*^*-/-*^ males and *Svep1*^*Δ/+*^*Apoe*^*-/-*^ females as breeders to generate experimental *Svep1*^*Δ/Δ*^*Myh11-CreER*^*T2*^*Apoe*^*-/-*^ (*Svep1*^*SMCΔ/Δ*^) and control *Myh11-CreER*^*T2*^*Apoe*^*-/-*^ (*Svep1*^*SMC+/+*^) mice. To activate Cre-recombinase, mice were injected intraperitoneally with 1 mg of tamoxifen (#T5648, Sigma-Aldrich) in 100 ml peanut oil (#P2144, Sigma-Aldrich) for 10 consecutive days starting at 6 weeks of age. Tamoxifen treatment was performed with all experimental and control mice in an identical manner. *Itgα9*^*flox/flox*^ (*Itgα9*^*Δ/Δ*^) mice were gifts from Dr. Dean Sheppard and Livingston Van De Water (Albany Medical College, New York), and *LysM-Cre* mice were provided from Dr. Babak Razani (Washington University School of Medicine, Saint Louis). We crossed *Itgα9*^*fl/fl*^ mice with *LysM-Cre* mice to generate *Itgα9*^*fl/fl*^*LysM-Cre* (*Itgα9*^*MACΔ/Δ*^) and control *Itgα9*^*+/+*^*LysM-Cre* (*Itgα9*^*MAC+/+*^) mice. All mice were housed in separate cages in a pathogen-free environment at Washington University School of Medicine animal facility and maintained on a 12 hr light/12 hr dark cycle with a room temperature of 22 ± 1°C.

### Diet and assessment of atherosclerosis

All experimental mice were fed a diet containing 21% fat and 0.2% cholesterol (#TD.88137, Envigo Teklad) for 8 and 16 weeks starting at 8 weeks of age. After HFD feeding, blood was collected from the retro-orbital plexus after 12 hr of fasting. Mice were euthanized by carbon dioxide inhalation. Plasma samples were prepared from the collected blood by centrifugation at 13,000 rpm for 10 min at 4°C. Total cholesterol (#STA-384), triglycerides (#STA-397), and glucose (#STA-681) in mouse plasma were determined using the appropriate kit (all purchased from Cell Biolabs, Inc). Hearts and whole aortas (from the aortic arch to the iliac artery) were harvested after perfusion with PBS. For *en face* analysis, isolated aortas were cleaned by removing perivascular fat tissues, opened longitudinally, and pinned onto black wax plates. After fixation with 4% paraformaldehyde overnight at 4°C, aortas were washed with PBS for 1 hr, and stained with 0.5% Oil Red O in propylene glycol (#O1516, Sigma-Aldrich) for 3 hr at room temperature. After staining, aortas were de-stained with 85% propylene glycol in distilled water for 5 min to reduce background staining and washed with distilled water for 15 min. For analysis of plaque in aortic root, hearts were fixed overnight with 4% paraformaldehyde at 4°C, washed with PBS for 1 hr, and embedded into OCT compound (#4583, Sakura^®^ Finetek). 5-µm-thick cryosections were stained overnight with 0.5% Oil Red O in propylene glycol, de-stained with 85% propylene glycol in distilled water for 5 min, and washed with distilled water for 15 min. Measurement of plaque was performed using 6-8 sections per artery to get the average value of size. The atherosclerotic plaque area was digitized and calculated using AxioVison (Carl Zeiss).

### Antibodies and reagents

To make the Svep1 protein, total RNA was purified from lung tissue of 8 week-old-mice by RNeasy^®^ kit (Life Technology). SuperScript IV First-Strand Synthesis System (Life Technology) with oligo d(T)_20_ primer was used to obtain full-length reverse transcripts. Then, double strand DNA was synthesized by PCR using PrimeSTAR GXL DNA Polymerase (Takara Bio) using forward 5’-ATGTGGTCGCGCCTGGCCTTTTGTTG and reverse 5’-AAGCCCGGCTCTCCTTTTCCTGGAACAATCAT primers, the amplicon was subcloned into pCMV6 plasmid (Origene Technology) in frame with Myc and Flag tag by In-Fusion HD EcoDry Cloning Plus System (Takara Bio). In order to insert a poly histidine tag for protein expression and purification, the Flag tag was replaced by a 10-histidine tag. Oligonucleotides 5’-CACCACCACCACCACCACCACCACCACCACTGAACGGCCGG and 5’-CCGTTCAGTGGTGGTGGTGGTGGTGGTGGTGGTGGTG coding for poly-histidine, stop codon and FseI compatible sticky end were synthetized, annealed and ligated to the vector, which was digested by EcoRV and FseI downstream of the myc tag and the stop codon respectively. The sequence of the full Svep1-myc-His_10_ construct was verified by Sanger sequencing. The protein was expressed in FreeStyle 293F cells (Invitrogen) grown in FreeStyle expression media. Transient transfection of the 3 µg ml^-1^ of vector DNA plus 9 µg ml^-1^ Polyethylenimine (PEI) (25 kDa linear PEI, Polysciences, Inc.) was used to express the protein in 2.5 × 10^6^ cells ml^-1^. After overnight incubation, the same volume of fresh media plus 1 mM (final concentration) valproic acid was added. The transfected cells were incubated in flasks in orbital shaker at 100 rpm in 37°C with 8% CO_2_ incubator for 48 hr, the conditioned media was collected by centrifugation at 300 rpm for 15 min. Cells were resuspended in fresh expression media and incubated for 48 hr. The conditioned media was collected, and this procedure was repeated one more time. Cell viability at the end of the experiment was >70%. The protein was purified in a NGC chromatographic system (BioRad Lab) in two steps: i) conditioned media was added to 1/10 of its volume of 10X PBS, and the pH check and fixed to 7.1. The protein was pulled down by a disposable column loaded with 5 ml Nuvia IMAC resin (BioRad Lab) at 1 ml min^-1^, washed with 50 ml PBS and the protein eluted in PBS plus 250 mM Imidazole, and ii) the eluted protein solution was concentrated 10 times in a Vivaspin concentrator 30 kDa cut off and loaded in a Superose 6 increase 10/300 (GE Life Sciences) with PBS as a carrier buffer. The fractions were evaluated by western blot probed with Myc tag antibody, and the purity of the protein was evaluated in PAGE-SDS 4-15% stained with Coomassie brilliant blue. More of the 95% of the protein in the gel corresponded to a single band. Further analysis by Mass Spectroscopy confirmed that more than 95% of the peptides detected corresponded to Svep1.

For immunofluorescent staining, anti-β-galactosidase (#ab9361, abcam, 1:1000), anti-Mac3 (#550292, clone M3/84, BD Biosciences, 1:100), anti-SMα-actin-cy3 (#C6198, clone 1A4, Sigma-Aldrich, 1:1000), anti-MCM-2 (#4007, Cell Signaling, 1:100) were used, and then visualized with anti-chicken-Alexa488 (#A11039), anti-rat-Alexa488 (#A21470), anti-rat-Alexa594 (#A21471, all purchased from Invitrogen, 1:400), and ProLong™ Gold antifade reagent with DAPI (#P36935, Invitrogen) were used. In case of detection of MCM-2 staining, samples were visualized with anti-rabbit-HRP (#7074S, Cell Signaling, 1:1000) followed by TSA^®^ Plus Cyanine 5 (#NEL745E001KT, PerkinElmer). For immunohistochemistry study, hematoxylin solution (#HHS80), eosin solution (#HT110180), Masson’s trichrome staining kit (#HT15-1KT, all purchased from Sigma-Aldrich), and Permount solution (#SP15-500, Fisher Chemicals) were used. For flow cytometry, following anti-mouse antibodies were used; anti-CD16/32 FcR blocker (#14-0161, eBioscience), PerCP-labeled anti-CD45 (#103129, clone 30-F11), BV510-labeled anti-CD11b (#101263, clone M1/70), BV421-labeled anti-CD64 (#139309, clone X54-5/7.1), PE/cy7-labeled anti-CD11c (#117317, clone N418), APC/cy7-labeled anti-MHCII (#107627, clone M5/114.15.2), FITC-labeled anti-F4/80 (#123108, clone BM8), BV605-labeled anti-CD19 (#115540, clone 6D5), APC-labeled anti-CD115 (#135510, clone AFS98), Alexa700-labeled anti-Ly6C (#128023, clone HK1.4), PE/Cy7-labeled anti-Ly6G (#127618, clone 1A8, all purchased from Biolegend), PE/cy5.5-labeled anti-CD4 (#35-0042-82, clone RM4-5, eBioscience), Alexa700-labeled anti-CD8a (#56-0081-80, clone 53-6.7, eBioscience), and PE-labeled anti-Itgα9β1 (#FAB3827P, R&D systems). Following anti-human antibodies were used. FcR blocker (#564219, BD biosciences), PerCP/cy5.5-labeled anti-CD45 (#368504, clone 2D1), Alexa700-labeled anti-CD3 (#300323, clone HIT3a), anti-CD19 (#115527, clone 6D5), anti-CD56 (#392417, clone QA17A16), FITC-labeled anti-CD15 (#301904, clone HI98), BV421-labeled CD66b (#305111, clone G10F5), APC/cy7-labeled anti-CD14 (#325619, clone HCD14), BV605-labeled anti-CD16 (#360727, clone B73.1), and PE-labeled anti-ITGα9β1 (#351606, clone Y9A2, all purchased from Biolegend). RBC lysis buffer (#423101, Biolegend), FoxP3 transcription factor staining buffer set (#00-5523, eBioscience), Leuko spin medium (#60-00091, Pluriselect) were used in flow cytometry experiments. Antibodies for western blotting include: Src (#2109), P-Src (#6943) FAK (#3285), P-FAK (#8556), Paxillin (#2542), P-Paxillin (#2541), Erk (#4695), P-Erk (#4370), p38 (#8690), P-p38 (#9211, all purchased from Cell signaling Technologies).

### Immunohistochemistry and immunofluorescent staining

For all immunohistochemistry and immunofluorescent study, we used 4% paraformaldehyde-fixed frozen sections with 5-µm-thickness. For immunofluorescent staining, slides were air-dried for 1 hr at room temperature and hydrated with PBS for 10 min. After permeabilization with 0.5% tritonX-100 for 10 min, sections were blocked with PBS containing 5% chicken serum (#S-3000, Vector Laboratories) with 0.5% tritonX-100 for 1 hr at room temperature. And then slides were incubated with the indicated antibodies. For hematoxylin and eosin (H&E) staining, air-dried slides were hydrated in PBS for 10 min, placed in hematoxylin solution for 10 min, and then rinsed in running tap water. After de-staining in 1% acetic acid for 5 min, slides were rinsed in tap water, and placed in 90% ethanol for 5 min. Slides were stained with eosin solution for 8 min, gradually dehydrated in ethanol solution (from 80% to 100%), and then incubated with xylene for 10 min followed by mounting with Permount solution. For Masson’s trichrome staining, air-dried slides were hydrated in distilled water for 10 min, placed in Mordant in Bousin’s solution for 1 hr at 56°C, and rinsed in running tap water for 5 min. After staining in hematoxylin solution for 10 min, slides were washed in running tap water for 10 min, rinsed in distilled water, placed in Biebrich scarlet-acid fuchsin solution for 15 min, and stained in aniline blue solution for 10 min. After rinsing in distilled water, slides were differentiated in 1% acetic acid for 3 min, gradually dehydrated in ethanol solution (from 80% to 100%), incubated with xylene for 10 min followed by mounting with Permount solution.

### RNAscope *in situ* hybridization (ISH)

For detection of *Svep1, Itgα9* RNA transcripts in both human and mouse artery tissues, a commercially available kit (#323100, RNAscope^®^ Multiplex Fluorescent Reagent Kit v2, Advanced Cell Diagnostics) was used according to the manufacturer’s instructions. Briefly, 4% paraformaldehyde-fixed mouse aortic root and human aortic wall, and LIMA frozen sections with 5-µm-thickness were air-dried for 1 hr at room temperature, and treated with hydrogen peroxide for 10 min to block endogenous peroxidase activity. After antigen retrieval by boiling in target antigen retrieval solution for 5 min at 95-100°C, slides were treated with protease III for 30 min at 40°C. Target probes (#406441, mouse *Svep1*; *#*540721, mouse *Itgα9*; #811671, human *SVEP1*; #811681, human *ITGA9*) were hybridized for 2 hr at 40°C, followed by a series of signal amplification and washing steps. Hybridization signals were detected by TSA^®^ Plus Cyanine 5, and co-stained with indicated antibodies. Slides were counterstained with DAPI by using ProLong™ Gold antifade reagent.

### Flow cytometry

For labeling mouse blood cells, blood was collected from the retro-orbital plexus, and red blood cells were removed using RBC lysis buffer (#00-4300-54, eBioscience). For labeling human blood cells, Leuko spin medium (Pluriselect) was used to isolate leukocytes from peripheral blood and buffy coat. In an experiment using mouse spleen, spleen cells were recovered from mice by cutting the spleen into small fragments, and incubated with 400 U collagenase D (#11-088-858, Roche applied science) for 30 min at 37°C. For labeling aortic single cell suspensions, isolated aortas were perfused with DPBS, and opened longitudinally. The whole artery was cut into 2–5 mm pieces, and incubated in a Hanks’ Balanced Salt Solution (HBSS) solution with calcium and magnesium containing 90 U ml^-1^ DNase I (#DN25), 675 U ml^-1^ collagenase I (#C0130), 187.5 U ml^-1^ collagenase XI (#C7657), and 90 U ml^-1^ hyaluronidase (#H1115000, all purchased from Sigma-Aldrich) for 70 min at 37 °C with gentle shaking. Non-specific binding to Fc receptors was blocked, and cells were incubated with the indicated antibodies for 30 min at 4°C. For intracellular staining, cells were fixed/permeabilized with the FoxP3 transcription factor staining buffer set. Flow cytometric analyses were performed using LSRFortessa™ instrument (BD Biosciences) and FlowJo software (Tree Star Inc).

### Bead labeling of Ly6C^low^ monocytes recruited into atherosclerotic plaque

After 8 weeks of HFD feeding, 200 µL of 1 µm Fluoresbrite yellow-green (YG) microspheres beads (#17154-10, Polysciences, Inc) diluted 1:4 in sterile DPBS were administered retro-orbitally. Labeling efficiency of blood monocytes was verified by flow cytometry 3 days after YG bead injection. Recruitment of YG-beads positive monocytes into plaque in aortic root was analyzed 1 day after checking labeling efficiency of YG beads. 5-µm-thick frozen sections of aortic root were stained with anti-Mac3, followed by anti-rat-Alexa594 antibody. And slides were mounted with ProLong™ Gold Antifade Mountant with DAPI. The number of YG-beads colocalized with Mac3 positive area was counted, or normalized with the percentages of labeled Ly6C^low^ monocytes.

### Aortic VSMC culture

Mouse aortic VSMCs were isolated from 8-week-old *Apoe*^*-/-*^ and *Svep1*^*+/-*^*Apoe*^*-/-*^, or same age of *Svep1*^*SMC+/+*^and *Svep1*^*SMCΔ/Δ*^ mice after tamoxifen injection for 10 consecutive days starting at 6 weeks of age. Briefly, thoracic aortas were harvested (3 mice per group were used), perivascular fat was removed, and then aortas were digested in 1 mg ml^-1^ collagenase II (#LS004174), 0.744 units ml^-1^ elastase (#LS002279), 1 mg ml^-1^ soybean trypsin inhibitor (#LS003570, all purchased from Worthington Biochemical Corporation), and 1% penicillin/streptomycin in HBSS for 10 min at 37°C with gentle shaking. After a pre-digestion with enzyme mixture, the adventitial layer was removed under the dissection microscope, and the intimal layer was removed by scrapping with forceps. Aortas were cut into small pieces, and completely digested in enzyme mixture at 37°C for 1 hr with gentle shaking. VSMCs were grown in 20% fetal bovine serum (FBS, Hyclone) containing Dulbecco modified Eagle medium/F12 (DMEM/F12) media (Gibco) with 100 U ml^-1^ penicillin/streptomycin at 37°C, 5% CO_2_ incubator. After 2 passages, VSMCs were changed to 10% serum. To stimulate VSMCs, 50 µg ml^-1^ human medium oxidized low density lipoprotein (oxLDL, #770202-7, Kalen biomedical) was used.

### Quantitative real time PCR

Gene expression was quantified by quantitative real time PCR. RNA was isolated using RNeasy® Mini Kit (#74106, Qiagen) according to manufacturer’s protocol, and QIAshredder homogenizer (#79656, Qiagen) to increase yield of quantification of RNA. cDNA was synthesized with High Capacity cDNA Reverse Transcript Kit (#4368814). Real time PCR was performed using both Taqman^®^ (#4444557) and SYBR™ Green (#A25742, all purchased from Applied Biosystems) assays. Ct values were normalized to *β-actin* (for Taqman^®^) and *gapdh* (for SYBR™ Green), and showed as expression relative to control. List of probes and primers used are in Supplemental Table 3.

### *In vitro* migration assay using peritoneal macrophage

*Itga9*^*MAC+/+*^ and *Itga9*^*MACΔ/Δ*^ mice were injected intraperitoneally with 1 ml 4% thioglycolate. After 5 days, the peritoneal cells were collected by lavage and placed in RPMI media containing 10% FBS for 60 min at 37°C. Non-adherent cells were removed after washing with PBS for 3 times, and adherent cells (more than 90% were peritoneal macrophages confirmed by flow cytometry) were placed in Trans-well inserts with a 5-µm porous membrane in a modified Boyden chamber. RPMI media containing 10% FBS with 50 or 200 ng ml^-1^ Svep1 protein was placed in the lower chamber. After allowing cell migration of 16 hr, inserts were removed from upper sider of the chamber, and nuclei of migrated cells to the lower side of the membrane were stained with DAPI. The number of migrated cells was determined by In Cell Analyzer 2000 (GE healthcare).

### Proliferation and adhesion assays

Wells of a 96 well plate were pre-coated with 30 µg ml^-1^ recombinant Svep1 protein or bovine serum albumin (BSA, as an inert protein control). Wells were subsequently blocked with 10 mg ml^-1^ BSA and washed twice with DPBS. Plates were ultraviolet (UV) sterilized before adding cells. For proliferation assays, primary VSMCs were collected and suspended in DMEM/F12 media containing 10% FBS. 2,000 cells were added to each well and incubated for 8 hr to assure complete cell adhesion. Media was then replaced with DMEM/F12 media containing 0.2% BSA and incubated for 12 hr to reduce basal proliferation rates. Cells were then incubated in BrdU dissolved in DMEM/F12 media containing 0.2% BSA for 30 hr. Predesigned Silencer Select siRNA constructs targeting *Itgα9* and negative control siRNA were obtained from ThermoFisher. Primary VSMCs were transfected using RNAiMAX transfection reagents according to the manufacturer’s protocol. Efficient *Itgα9* knockdown was confirmed by qPCR. Cells were trypsinized 24 hr after transfection and used for the proliferation assay. DAPT or DMSO (carrier) were added to cells throughout the indicated experiment at a concentration of 25 µM. For using peritoneal macrophages, 4% thioglycolate-elicited peritoneal macrophages from *Itgα9*^*MAC+/+*^ and *Itgα9*^*MACΔ/Δ*^ mice were suspended in BrdU-containing RPMI 1640 media that also contained 10% FBS. 25,000 cells were added to each well since peritoneal macrophages have lower proliferation rates than VSMCs in culture. 50 µg ml^-1^ oxLDL was added to the indicated cells at the beginning of this incubation. An ELISA for incorporated BrdU was then performed using kit instructions (#6813, Cell Signaling Technologies) after incubation for 36 hr. Adhesion assays were performed in precoated 96-well plates blocked with 100 mg ml^-1^ BSA. Blocking conditions were empirically derived to minimize non-specific cell adhesion. After a 5 (for THP-1 cells) or 15 (for VSMCs) min incubation, non-adhered cells were removed by gently centrifuging the plates upside down. VSMCs were counted manually and THP-1 cells were counted by automated microscopy after staining cells with DAPI.

### Western blotting assay

Cells were resuspended in serum free media (SFM), and incubated with gentle agitation to prevent cell attachment and reduce basal signaling. Cells were washed with SFM then seeded on BSA-blocked plates coated with either BSA, VCAM-1, Svep1, or Svep1^CADrv^. Concentrations of VCAM-1 and Svep1 were derived empirically to prevent signal saturation. BSA concentrations always matched the Svep1 concentration. It is only appropriate, therefore, to compare between BSA, Svep1, and Svep1^CADrv^ groups. Cells were briefly centrifuged to the bottom of the wells and incubated for 8 min (for VSMCs) or 15 min (for THP-1 cells) before lysis with cell lysis buffer (#9803, Cell Signaling Technologies) containing a cocktail of protease and phosphatase inhibitors. Western blots were performed by standard techniques, as briefly follows. Protein content was determined using a bicinchoninic acid assay with BSA standards (#23225, Pierce™ BCA Protein Assay Kit). Cell lysates were then reduced with DTT in lithium dodecyl sulfate sample buffer (#NP0007, Invitrogen). Equal protein amounts were added to polyacrylamide gels (#4561086, BioRad) and electrophoresed prior to transferring to a nitrocellulose or polyvinylidene fluoride membrane (#1620260, BioRad). Membranes were blocked in 5% BSA/Tris-Buffered Saline with tween 20 for 30 min. The indicated primary antibodies were incubated with the pre-blocked membranes for overnight at 4°C. Membranes were washed with Tris-Buffered Saline with tween 20, probed with fluorescent secondary antibodies, and imaged. β-actin or β-tubulin served as a loading control.

### Bulk RNA sequencing and analysis

Primary VSMCs were plated on wells precoated with 30 µg ml^-1^ recombinant Svep1, Svep1^CADrv^ protein or BSA (as an inert protein control). Wells were subsequently blocked with 10 mg ml^-1^ BSA and washed twice with DPBS. Plates were UV sterilized before adding cells. Primary VSMCs were collected, resuspended in DMEM/F12 media containing 10% FBS, plated on precoated, blocked wells, and incubated for 8 hr to ensure complete cell adhesion. Media was replaced with fresh DMEM/F12 containing 1% FBS and incubated for 12 hr before collection. RNA was collected using RNeasy® Mini Kit (#74106, Qiagen). Atherosclerotic aortic arches (including the aortic root, arch, and the proximal regions of its branching vessels) from mice were used as the source of RNA for the later RNAseq experiment. These tissues were isolated and separated from the perivascular adipose prior to storing in RNAlater (#AM7021, Thermofisher) prior to total RNA extraction using nucleoZOL (Macherey-Nagel). cDNA for validation was synthesized with High Capacity cDNA Reverse Transcript Kit (#4368814, Applied Biosystems), following standard protocols.

Samples were prepared according to library kit manufacturer’s protocol, indexed, pooled, and sequenced on an Illumina HiSeq. Basecalls and demultiplexing were performed with Illumina’s bcl2fastq software and a custom python demultiplexing program with a maximum of one mismatch in the indexing read. RNA-seq reads were then aligned to the Ensembl release 76 primary assembly with STAR version 2.5.1a (Dobin et al., 2013). Gene counts were derived from the number of uniquely aligned unambiguous reads by Subread:featureCount version 1.4.6-p5 (Liao et al., 2014). Isoform expression of known Ensembl transcripts were estimated with Salmon version 0.8.2 (Patro et al., 2017). Sequencing performance was assessed for the total number of aligned reads, total number of uniquely aligned reads, and features detected. The ribosomal fraction, known junction saturation, and read distribution over known gene models were quantified with RSeQC version 2.6.2 (Wang et al., 2012). All gene counts were then imported into the R/Bioconductor package EdgeR (Robinson et al., 2010) and TMM normalization size factors were calculated to adjust for samples for differences in library size. Ribosomal genes and genes not expressed in the smallest group size minus one sample greater than one count-per-million were excluded from further analysis. The TMM size factors and the matrix of counts were then imported into the R/Bioconductor package Limma (Ritchie et al., 2015). Weighted likelihoods based on the observed mean-variance relationship of every gene and sample were then calculated for all samples with the voomWithQualityWeights (Liu et al., 2015). The performance of all genes was assessed with plots of the residual standard deviation of every gene to their average log-count with a robustly fitted trend line of the residuals. Differential expression analysis was then performed to analyze for differences between conditions and the results were filtered for only those genes with Benjamini-Hochberg false-discovery rate adjusted *P*-values less than or equal to 0.05. One sample in the aortic arch experiment was independently identified as an outlier by standard quality control methods. This group was excluded from downstream analyses.

For each contrast extracted with Limma, global perturbations in known Gene Ontology (GO) terms, KEGG pathways, and InterPro domains were detected using the Database for Annotation, Visualization and Integrated Discovery (DAVID) (Huang da et al., 2009) on significantly dysregulated transcripts or using the R/Bioconductor package GAGE (Luo et al., 2009) to test for changes in expression of the reported log 2 fold-changes reported by Limma in each term versus the background log 2 fold-changes of all genes found outside the respective term. The R/Bioconductor package heatmap3 (Zhao et al., 2014) was used to display heatmaps across groups of samples for each GO or MSigDb term with a Benjamini-Hochberg false-discovery rate adjusted p-value less than or equal to 0.05. Perturbed KEGG pathways where the observed log 2 fold-changes of genes within the term were significantly perturbed in any direction compared to other genes within a given term with *P*-values less than or equal to 0.05 were rendered as annotated KEGG graphs with the R/Bioconductor package Pathview (Luo and Brouwer, 2013).

### Notch signaling assays

Cells were collected, resuspended in SFM, and incubated for 1 hr with gentle agitation before seeding on tissue culture wells that were precoated and blocked, as described in previous sections. Cells were collected for analysis after 4 hr of growth on the indicated substrate. *Svep1*^*SMC+/+*^and *Svep1*^*SMCΔ/Δ*^ VSMCs were collected for analysis after 72 hr of incubation in SFM to obtain basal Notch signaling. Notch target gene primers used for the qPCR are listed in Supplementary Table 3.

### Analysis of cytokine and chemokine biomarkers

MILLIPLEX MAP Mouse Cytokine/Chemokine Magnetic Bead Panel-Immunology Multiplex Assay (#MCYTOMAG-70K), MILLIPLEX MAP Mouse Angiogenesis/Growth Factor Magnetic Bead Panel-Cancer Multiplex Assay (#MAGPMAG-24K), and MILLIPLEX MAP Mouse Cardiovascular Disease (CVD) Magnetic Bead Panel 1-Cardiovascular Disease Multiplex Assay (#MCVD1MAG-77K-02, all from Millipore Sigma) were used to analyze cytokines and chemokines from mouse plasma. All kits were used according to manufacturer recommended protocols. Briefly, the Luminex FLEXMAP 3D® (Luminex Corporation, Austin, TX) instrument was used to sort the magnetic polystyrene beads and measure the phycoerythrin (PE) tagged detection antibody signal. Fifty beads from each analyte were measured. The median fluorescent intensity (MFI) was compared against the standard curve to calculate the pg ml^-1^ or ng ml^-1^ using Milliplex Analyst 5.1 software (VigeneTech.com) and a 5-parameter logistic curve fit algorithm.

### BMDM isolation and culture

6-to 8-week-old *Itgα9*^*MAC+/+*^ and *Itgα9*^*MACΔ/Δ*^ mice were euthanized by carbon dioxide inhalation, and soaked in 75% ethanol. Then, femurs and tibias were harvested and bone-marrow cells were obtained by flushing bones and differentiated for 7 days in DMEM media supplemented with 50 mg ml^-1^ recombinant macrophage-colony stimulating factor (M-CSF, R&D systems), 20% heat-inactivated FBS, and antibiotics.

### Mendelian Randomization

To estimate the causal effect of SVEP1 plasma protein levels on risk of CAD, hypertension, and type 2 diabetes (T2D), we performed Mendelian Randomization using summary statistics from publicly available datasets. Genome-wide summary statistics for risk of CAD were obtained from a meta-analysis of CAD using data from CARDIoGRAMPlusC4D and the UK Biobank as previously described (van der Harst and Verweij, 2018). Genome-wide summary statistics for hypertension and T2D were obtained from the IEU GWAS database (Hemani et al., 2018) using association results from the UK Biobank. Summary statistics for primary hypertension (ICD 10 code I10) as a secondary diagnosis (IEU GWAS ID “ukb-b-12493”) were used for hypertension while summary statistics for diabetes diagnosed by a doctor (IEU GWAS ID “ukb-b-10753”) were used for T2D.

A genome-wide association study to identify protein quantitative trait loci using a SomaLogic aptamer-based protein assay has previously been described (Sun et al., 2018). Two aptamers (SVEP1.11109.56.3 and SVEP1.11178.21.3) were used to estimate SVEP1 protein concentration. We obtained genome-wide summary statistics for both aptamers which produced highly similar results; for simplicity, results from the analysis using the SVEP1.11178.21.3 aptamer were reported. As trans-pQTLs might affect protein levels in a variety of manners, we focused our analysis on cis-pQTLs by only including variants in a 1Mb window surrounding SVEP1 which associated with plasma SVEP1 concentration at a level exceeding genome-wide significance (*P*-value for SVEP1 concentration < 5 × 10^−8^). We filtered these SNPs using pair-wise linkage disequilibrium estimated from the 1000 Genomes Project European samples in order to obtain an independent (r^2^ < 0.3) set of SNPs for the causal analysis. Causal estimates were calculated using the inverse-variant weighted method implemented in the R package TwoSampleMR (Hemani et al., 2017; Hemani et al., 2018).

### Statistical analysis

For animal model data, a two-group independent t-test, one-way analysis of variance (ANOVA), or two-way ANOVA were used, provided the data satisfied the Shapiro-Wilk normality test. Otherwise, the Mann-Whitney U test, Kruskal-Wallis one-way ANOVA test, and Friedman two-way ANOVA test were used. Bonferroni correction was used for post-hoc multiple comparison in ANOVA. Unless otherwise stated, cellular assays were analyzed by an unpaired, two-tailed t-test. Statistical analyses were performed with GraphPad Prism.

