## Supplementary Material for "SVEP1, a novel human coronary artery disease locus, promotes atherosclerosis"

**Figure S1.** ***SVEP1*** **expression in health and disease.**

(**A**) Abbreviated list of GTEx tissues in order of *SVEP1* expression levels. (**B**) *SVEP1* expression in atherosclerotic carotid arterial tissue, relative to adjacent, paired arterial tissue. Red line represents the reference tissue. Data collected from GEO GSE43292. Paired t-test. (**C**) *SVEP1* expression in medial and intimal explants from control and diabetic patients. Data collected from GEO GSE13760. Two-tailed t-test. (**D**) *Svep1* expression in relevant cell types of the murine aorta as previously published (Kalluri et al., 2019). (**E**) Quantification of *Svep1* fluorescence intensity in atherosclerotic plaque. (**F**) Quantification of *Svep1* fluorescence intensity in *Svep1*$^{SMC\Delta/\Delta}$ animals. Mann-Whitney test for (E and D) *$P < 0.05$; ***$P < 0.001$.

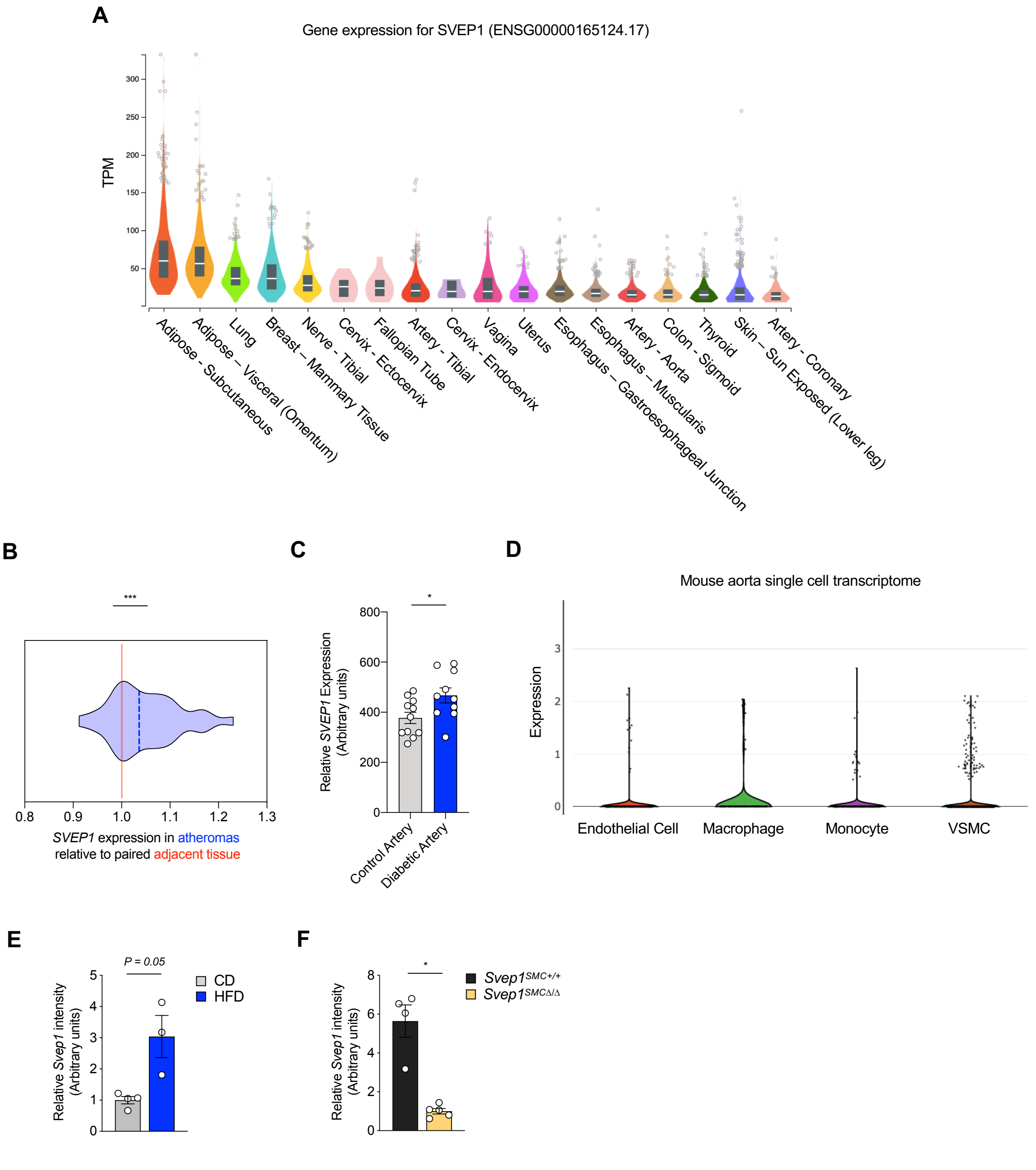

**Figure S2. Whole body knockout of *Svep1* does not significantly alter plaque composition after 8 weeks of HFD feeding.**

(**A**) SMα-actin staining *n* =12/group. Scale bar, 500 µm. (**B**) Collagen staining using by Masson's trichrome stain, *n* = 11-12/group. (**C**) Necrotic core outlined on H&E-stained tissues. *n* =12/group. Scale bars in B and C, 200 µm. L, lumen. All data were analyzed with unpaired nonparametric Mann-Whitney test, and shown as the mean ± SEM. *$P < 0.05$; *NS*, not significant.

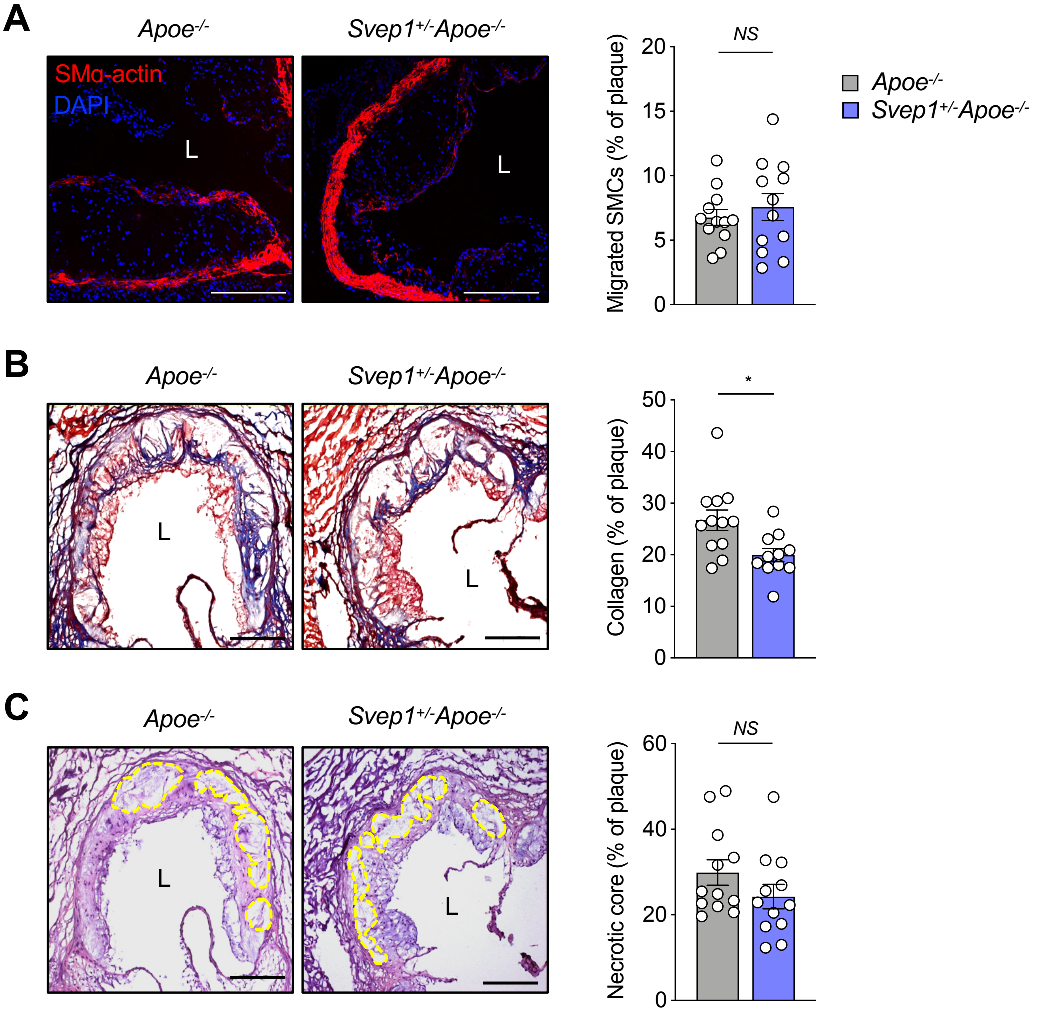

**Figure S3. VSMC-specific *Svep1* deficiency reduces atherogenesis and plaque complexity, continued.**

(**A**) Mac3 staining of aortic root. (**B**) Collagen staining of aortic root by Masson's trichrome stain. (**C**) Necrotic core outlined on H&E-stained sections. *n* = 13-15/group (A through C). (**D**) Body weight of $Svep1^{SMC+/+}$ and $Svep1^{SMC\Delta/\Delta}$ mice during HFD feeding. (**E**) Plasma total cholesterol, triglycerides, and glucose. (**F**) *En face* Oil Red O-stained aortas. Outlined areas indicate the aortic arch regions magnified in left panels. Quantification of Oil Red O-stained area in each aortic arch and whole artery. (**G**) Oil Red O-stained aortic root cross-sections. Quantification of Oil Red O-stained area. n=8-9/group. Scale bars, 50 µm (in A), 200 µm (in B and C), 500 µm (in G). 8 weeks of HFD feeding (A through C), and 16 weeks of HFD feeding (D through G). Data were analyzed with One-way ANOVA test (D), unpaired nonparametric Mann-Whitney test (A through C, E through G), and shown as the mean ± SEM. *$P < 0.05$; **$P < 0.01$; *NS*, not significant.

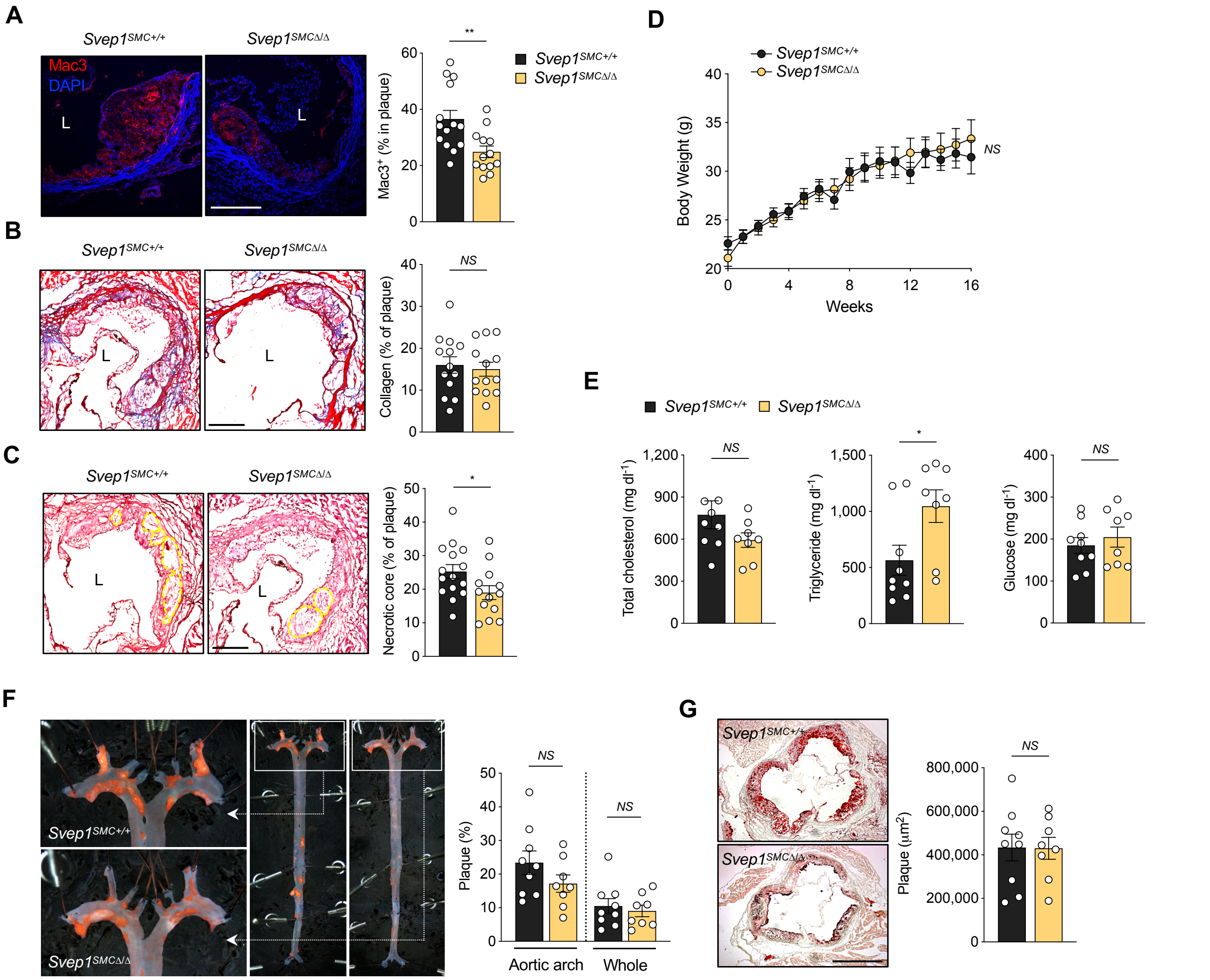

**Figure S4. The effect of the CAD-associated SVEP1 D2702G missense polymorphism and SVEP1 protein levels in humans and mice.**

(**A**) Relative mRNA levels by genotype as measured in the GTEx project. There is no significant difference in *SVEP1* expression by *SVEP1* D2702 genotype in coronary artery ($P = 1.0$), aorta ($P = 0.32$), or cultured fibroblasts ($P = 0.9$), with the latter being included as it is the highest expressing cell sample in GTEx. (**B**) The effect of each chromosome 9 variant on plasma SVEP1 protein level as estimated by the SVEP1.11109.56.3 and SVEP1.11178.21.3 aptamers. (**C**, **D**) The respective leftward panels indicate the estimated effect (with 95% confidence intervals) of each variant included in the Mendelian Randomization analysis on plasma SVEP1 level and risk of hypertension (HTN, C) or type 2 diabetes (T2D, D). The red line indicates the causal effect estimate ($P = 2 \times 10^{-15}$ for HTN; $P = 0.0004$ for T2D). The respective rightward panels indicate the estimated causal effect (with 95% confidence intervals) of each SNP included in the Mendelian Randomization analysis for a one unit increase in SVEP1 level. These are plotted along with the overall summary estimate from the causal analysis. (**E**) Body weight of $Apoe^{-/-}$ and $Svep1^{G/G}Apoe^{-/-}$ mice during 8 weeks of HFD. (**F**) Total plasma cholesterol, triglyceride and glucose after 8 weeks of HFD. (**G**) Body weight of $Apoe^{-/-}$ and $Svep1^{G/G}Apoe^{-/-}$ mice during 16 weeks HFD. (**H**) Total plasma cholesterol, triglyceride and glucose after 16 weeks of HFD. (**I**) Oil Red O-stained aortic root cross-sections of mice after 8 weeks HFD. Quantification of Oil Red O-stained area. (**J**) Oil Red O-stained aortic root cross-sections of mice after 16 weeks HFD. Quantification of Oil Red O-stained area. Scale bar, 500 µm (in G and H). *NS*, not significant.

**Figure S4. The effect of the CAD-associated SVEP1 D2702G missense polymorphism and SVEP1 protein levels in humans and mice.**

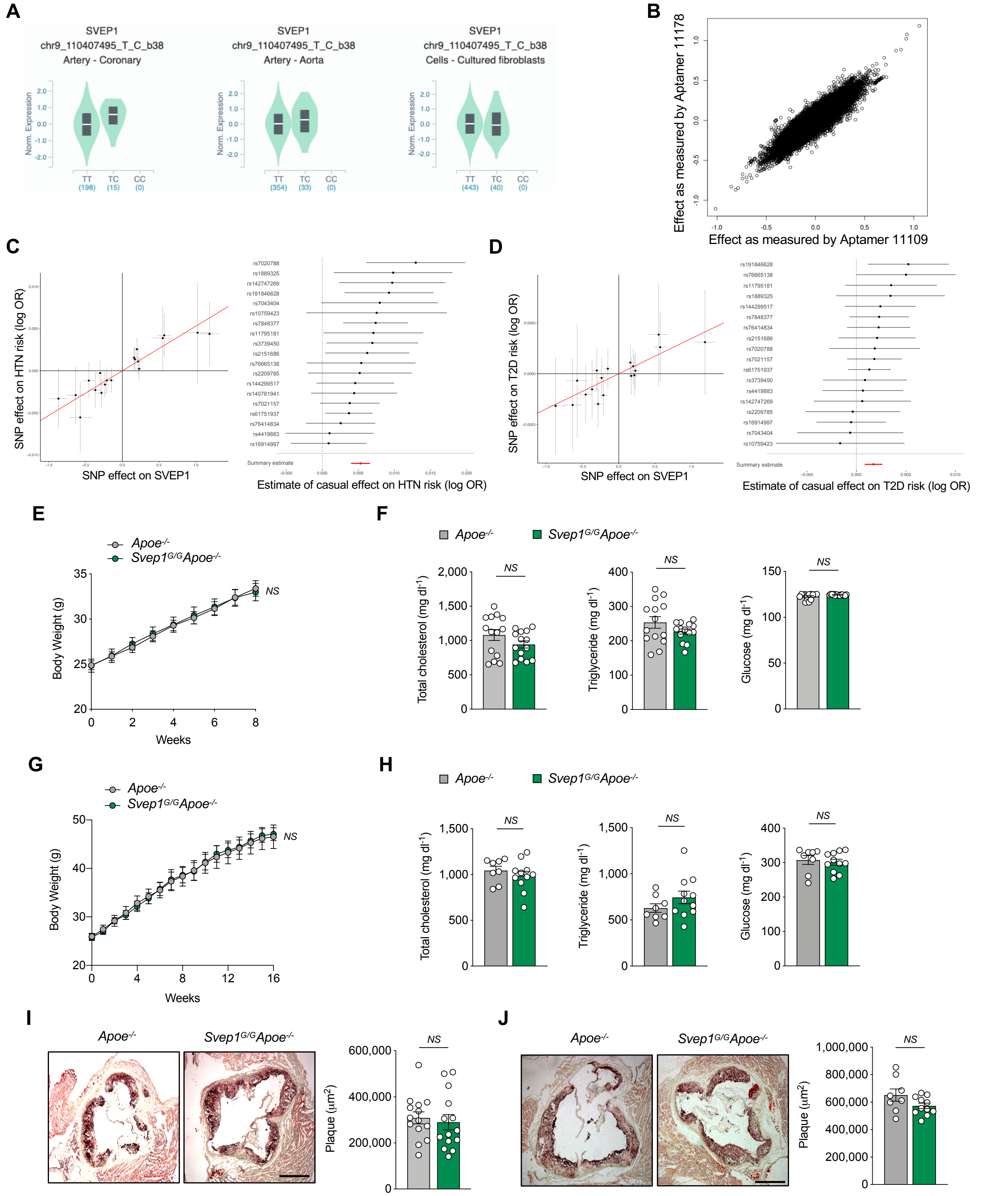

**Figure S5. *Itga9* is expressed in both VSMCs and macrophages in mouse arteries.**

(**A**) Abbreviated list of GTEx tissues in order of *ITGA9* expression levels. (**B**) *Itgα*9 expression in the murine aorta. Data collected from (Kalluri et al., 2019) (**C**) Proliferation of $Itga9^{MAC+/+}$ and $Itga9^{MAC\Delta/\Delta}$ macrophages. Cells were grown on with immobilized Svep1 or BSA for 8 hr and then treated with 50 µg ml$^{-1}$ oxLDL in the culture media for 36 hr. Incorporated BrdU was measured by ELISA. Data were analyzed with One-way ANOVA test, and shown as the mean ± SEM. *NS*, not significant. (**D**) Immunoblots of proximal integrin signaling kinases and downstream p38 from VSMCs adhered to control, or Svep1, or Svep1$^{CADrv}$-treated plates. β-actin was used as loading control.

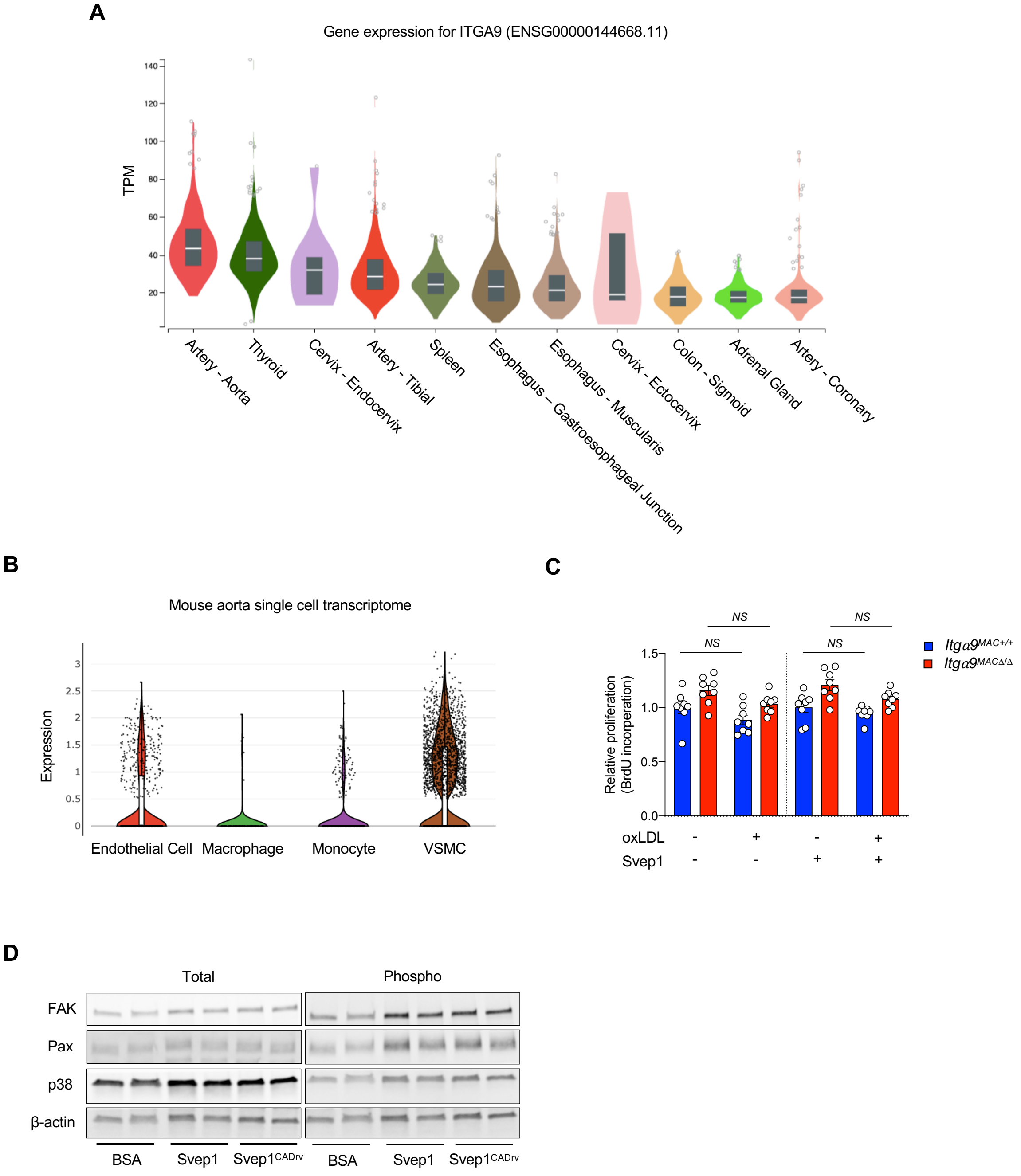

**Figure S6. Svep1 promotes local inflammation in atherosclerosis**

(**A**) qPCR determination of *Ccl2*, *Spp1*, and *Cxcl5* expression in aortic arches. Each gene was normalized to *Gapdh.* Two-tailed t-test, and shown as the mean ± SEM. (**B**) Analysis of cytokine and chemokine biomarkers from 8 weeks HFD-fed *Svep1*$^{SMC+/+}$ and *Svep1*$^{SMC\Delta/\Delta}$ mice. sICAM-1, soluble intercellular adhesion molecule-1; P-selectin, soluble P-selectin; Ang-2, angiotensin-2; IL-6, interleukin-6; KC, C-X-C motif ligand1 (CXCL1); *NS*, not significant.

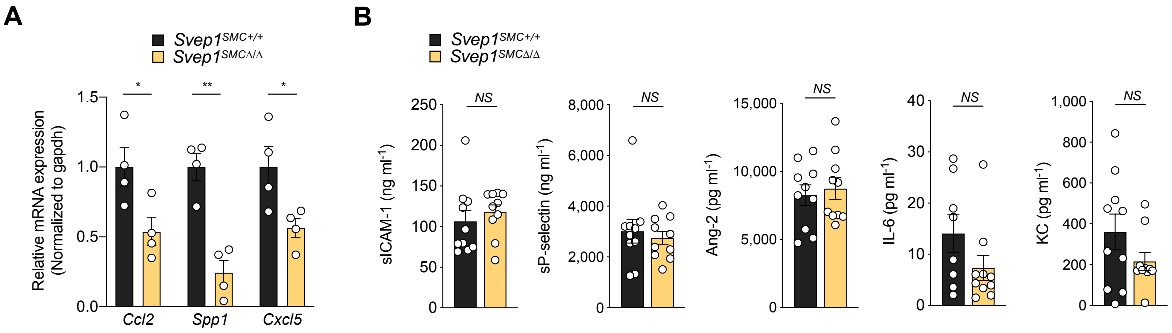

**Figure S7. Svep1 promotes local inflammation in atherosclerosis**

(**A**) ITGA9 expression in neutrophils and monocyte subsets ($CD14^{low}CD16^{+}$ non-classical, $CD14^{high}CD16^{+}$ intermediate, and $CD14^{+}CD16^{-}$ classical monocytes) in human blood. (**B**) *ITGA9* expression in human aortic wall and LIMA cross-sections from patients using ISH. Tissues were co-stained for CD68. (**C**) Gating strategy and histogram for Itgα9β1 expression in mouse blood neutrophils ($CD11b^{+}Ly6G^{+}$), $Ly6C^{low}$ ($CD11b^{+}Ly6C^{low}$), and $Ly6C^{high}$ ($CD11b^{+}Ly6C^{high}$) monocytes. (**D**) Gating strategy and histogram of Itgα9β1 expression in the subpopulations. Splenocytes isolated from *Apoe*$^{-/-}$ mice were incubated with 400 U $ml^{-1}$ collagenase D for 30 min at 37°C followed by incubation with 25 μg $ml^{-1}$ oxLDL or vehicle control (PBS) for 48 hr. Gating strategy for detection of macrophages, DCs, neutrophils, and $Ly6C^{high}$ monocytes. Briefly, $CD45^{+}$ cells were detected after removal of auto-fluorescence with unused fluorescence, for further analysis of $CD64^{+}F4/80^{+}$ macrophages. Among $CD64^{-}F4/80^{-}$ cells, $MHCII^{high}CD11c^{+}$ DCs were detected following detection of $CD11b^{+}Ly6G^{+}$ neutrophils and $CD11b^{+}Ly6C^{high}$ monocytes from $CD11b^{+}$ cells with Ly6G and Ly6C expression. Macrophages ($CD64^{+}CD11b^{+}$), DCs ($MHCII^{high}CD11c^{+}$), neutrophils ($CD11b^{+}Ly6G^{+}$), and $Ly6C^{high}$ ($CD11b^{+}Ly6C^{high}$) monocytes from mice group. (**E**) Gating strategy for detection of B lymphocytes and T lymphocytes including cytotoxic $CD8^{+}$ ($CD8^{+}$ $T_{cyt}$), regulatory $CD8^{+}$ ($CD8^{+}$ $T_{reg}$), effector $CD4^{+}$ ($CD4^{+}$ $T_{eff}$), and regulatory $CD4^{+}$ T cells ($CD4^{+}$ $T_{reg}$). Histograms of Svep1 and Itgα9β1 expression in B and T lymphocytes. (**F**) Adhesion of THP-1 cells to increasing concentrations of Svep1. Data are normalized to BSA control. Nonadherent cells were removed by centrifugation. (**G**) THP-1 cells were seeded on wells containing the indicated protein. Cells were lysed and protein lysates were subject to immunoblotting with the indicated antibody. BSA served as a negative control and VCAM-1 served as positive control. β-tubulin was used as loading control. (**H**) *In vivo* monocyte recruitment assay. YG-beads were administered to mice retro-orbitally after 8 weeks of HFD feeding. Labeling efficiency of $Ly6C^{low}$ monocytes was verified and then YG-beads in the aortic root plaque were subsequently analyzed. Gating strategy for detection of YG-bead labeling by $Ly6C^{low}$ monocytes. Among $CD11b^{+}$ cells, Ly6G positive neutrophils ($CD11b^{+}CD115^{-}Ly6G^{+}$) were gated out, following further analysis of $CD115^{+}$ cells (total monocytes) with Ly6C expression. YG-beads were only detected in $Ly6C^{low}$ not $Ly6C^{high}$ monocytes. (**I**) Labeling efficiency as the percentage and total number of YG-bead positive $CD115^{+}$, $Ly6C^{low}$, and $Ly6C^{high}$ monocytes. *NS*, not significant.

### Figure S7. Svep1 promotes local inflammation in atherosclerosis

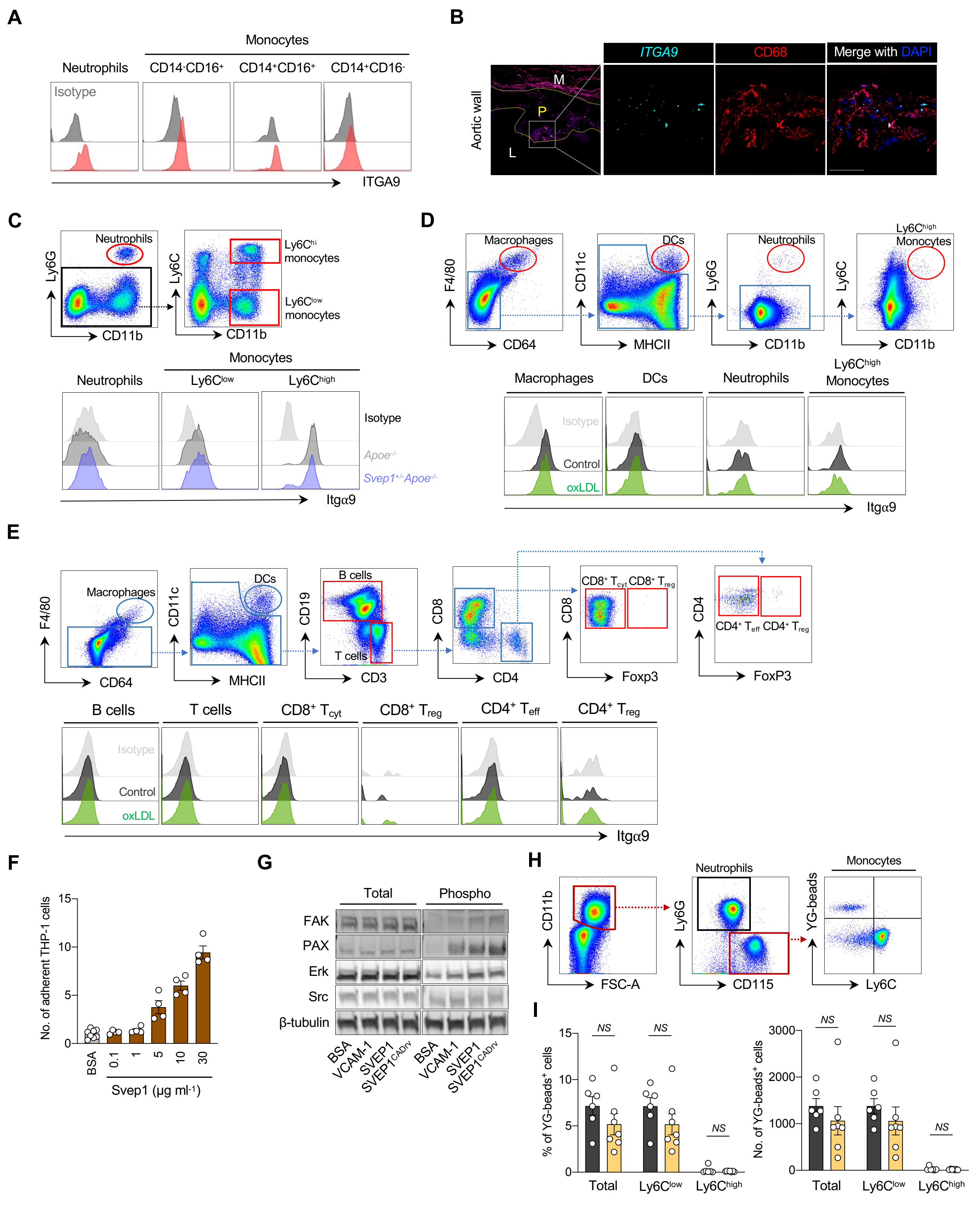

**Table S1. VSMC response to Svep1 variants.** Transcripts detected and gene ontology enrichment analysis in RNA collected from primary VSMCs exposed to BSA, Svep1, or Svep1$^{CADrv}$ by RNAseq analysis. See SVEP1-Jung-Supplementary-Table1.xlsx.

**Table S2. Gene ontology enrichment of atherosclerotic aortic arches.** Transcripts detected and gene ontology enrichment analysis of RNAseq from atherosclerotic aortic arches of HFD-fed *Svep1*$^{SMC+/+}$ and *Svep1*$^{SMC\Delta/\Delta}$ mice. See SVEP1-Jung-Supplementary-Table2.xlsx.

**Table S3. Primer Table.** List of oligonucleotides used for qPCR studies.

| **Taqman®** | **Catalog No.** | **Company** |
|---|---|---|
| *Svep1* | Mm00465702_m1 | ThermoFisher Scientific |
| *CD36* | Mm00432403_m1 | ThermoFisher Scientific |
| *Itgα9* | Mm00519317_m1 | ThermoFisher Scientific |
| *β-actin* | Mm02619580_m1 | ThermoFisher Scientific |
| **SYBR™ Green** | **Forward primer** | **Reverse primer** |
| *Ccl2* | CAAGAAGGAATGGGTCCAGA | GCTGAAGACCTTAGGGCAGA |
| *Spp1* | TCACCATTCGGATGAGTCTG | ACTTGTGGCTCTGATGTTCC |
| *Cxcl5* | ATCACCTCCAAATTAGCGATCA | TTCTGTTGCTGTTCACGCT |
| *SMα-actin* | GCATCCACGAAACCACCTA | CACGAGTAACAAATCAAAGC |
| *Myh11* | CATGGACCCGCTAAATGACA | CAATGCGGTCCACATCCTTC |
| *Cxcl1* | GCTTGAAGGTGTTGCCCTCAG | AAGCCTCGCGACCATTCTTG |
| *Il-6* | AGTTGCCTTCTTGGGACTGA | TCCACGATTTCCCAGAGAAC |
| *Hey1* | GAAGCGCCGACGAGACCGAATCAA | CAGGGCGTGCGCGTCAAAATAACC |
| *Hey2* | CGACGTGGGGAGCGAGAACAAT | GGCAAGAGCATGGGCATCAAAGTA |
| *Heyl* | AGACCGCATCAACAGCA | CAAGTGATCCACGGTCAT |
| *Hes1* | GACGGCCAATTTGCTTTC | GACACTGCGTTAGGACCC |
| *Gapdh* | TCACCACCATGGAGAAGGC | GCTAAGCAGTTGGTGGTGCA |
